# Specialization between foliar fungal endophyte and plant communities decreases toward the tropics

**DOI:** 10.64898/2026.10.07.756763

**Authors:** Austen Apigo, Rodolfo Salas-Lizana, José Rubén Montes, Rafael Pinheiro, Luis C. Mejía, Ryoko Oono

## Abstract

Associations among species are hypothesized to become fewer and more specialized toward the tropics. However, evidence for the latitudinal specialization gradient (LSG) remains mixed and restricted to plant-animal and host-parasite models, with increasing recognition that our understanding of LSG is often complicated by differences in species richness and sampling effects. Here, we tested whether specialization between foliar fungal endophytes—fungi that live within plant leaves—and their plant hosts increased toward the tropics by surveying 1,539 plants that spanned a 55-degree latitudinal gradient in Central and North America. Contrary to expectations, endophytes and plants formed a greater number of associations that were more evenly distributed toward tropical latitudes and were thus less specialized, even after accounting for differences in endophyte and plant community structure. Endophyte-plant communities were most specialized where evaporative demand and plant richness were low, and where water availability and primary productivity were high. Our findings show a reversed LSG where the specialization between endophytes and plants decreased toward the tropics and provides evidence that many species of endophytes can coexist even when there is high overlap in their host niches.

## Introduction

Biologists have long sought to understand the latitudinal diversity gradient, with a focus on the climax of species richness in the tropics (Rohde 1992, Gaston 2000, Hillebrand 2004, Kreft and Jetz 2007). The latitudinal niche-breadth hypothesis helps explain this by suggesting that low seasonality and high productivity in the tropics stabilize populations, allowing species to evolve narrower ecological niches and coexist in greater numbers (MacArthur 1972, Vázquez and Stevens 2004). This led to the proposal of a latitudinal specialization gradient (LSG), which predicts that species associate with smaller proportions of the community, increasing biotic specialization, toward the tropics (MacArthur 1972, Moles and Ollerton 2016, Dyer and Forister 2019). In turn, greater biotic specialization is thought to explain why species richness increases toward the tropics (Dobzhansky 1950, MacArthur 1965, Moles and Ollerton 2016, Dyer and Forister 2019). However, evidence for increasing specialization toward the tropics is mixed among plant and animal communities (Ollerton and Cranmer 2002, Olesen and Jordano 2002, Dyer et al. 2007, Dalsgaard et al. 2011, 2017, Schleuning et al. 2012, Trøjelsgaard and Olesen 2013, Morris et al. 2014, Forister et al. 2015, Dáttilo and Vasconcelos 2019, Sonne et al. 2020, Doré et al. 2021, Nakadai et al. 2021, Luna et al. 2022, Pinheiro et al. 2023), challenging the generality of the LSG and its connection to the latitudinal diversity gradient.

The latitudinal niche-breadth hypothesis predicts a positive correlation between specialization and species richness. However, a greater number of species in a community is also thought to create more ecological opportunities for species associations, which might expand available resources and reduce competitive interactions (MacArthur 1965, Schemske et al. 2009). This could result in wider, overlapping species niches and decreased specialization in places with greater species richness such as the tropics (MacArthur 1965, Pianka 1974, Cirtwill et al. 2015, Dehling et al. 2022). For example, studies of plant-pollinator and plant-seed disperser communities have shown decreasing specialization toward the tropics in response to increasing plant diversity (Schleuning et al. 2012, Dalsgaard et al. 2017). Therefore, examining whether species richness drives specialization, rather than the reverse, may help resolve the lack of consensus surrounding LSG.

Indeed, meta-analyses of plant and animal communities have shown that network size (the number of species in a community) explains more variation in specialization than climate, primary productivity, or latitude at global scales (Morris et al. 2014, Doré et al. 2021, Brimacombe et al. 2022a, 2022b). A further challenge in assessing the relationship between species richness and specialization is that networks of greater size become more specialized simply because the potential number of species associations increases exponentially, while the number of observed associations often do not (Blüthgen et al. 2006). Thus, increased measures of specialization could be due to increasing species richness, greater sampling effort, or a combination of both. Because the information needed to account for differences in sampling methods and scale is inconsistently available among public data (Brimacombe et al. 2022b), meta-analyses often include network size in statistical models to control for biases rather than to test for an effect of species richness. Sampling biological communities in a consistent manner across different ecosystems remains a rare practice (Dyer and Forister 2019, Brimacombe et al. 2022b) but is critical to test the relationship between species richness and specialization.

Plant-animal and host-parasite systems have been widely used as models to test for LSG, but whether LSG explains the biogeography of plant-associated microbial communities is still an open question. Many fungi form localized, asymptomatic infections within the interiors of photosynthetic tissues as foliar fungal endophytes (hereafter, endophytes; Schulz and Boyle 2005, Arnold 2007). Their cryptic nature and ubiquity across all major lineages of land plants have motivated work to understand where endophytes are most diverse, with current evidence agreeing that tropical rainforests support the greatest richness of endophytes (Arnold and Lutzoni 2007; this study). According to the latitudinal niche-breadth hypothesis, endophyte-plant specialization should also be greatest in the tropics. However, studies suggest generalist endophyte species dominate tropical plant communities (Arnold and Lutzoni 2007, Higgins et al. 2011, 2014), while endophyte communities have been shown to be highly distinct in species composition among boreal plant lineages (U’Ren et al. 2019), suggestive of a reversed LSG. Despite these findings, a test of LSG has yet to be conducted with a consistent sampling of temperate and tropical endophyte communities. To address this gap, we systematically sampled communities of endophytes and plants across 20 forested sites spanning a 55-degree latitudinal gradient in Central and North America (Figure 1, Tables S1-S2). We applied amplicon sequencing to characterize endophyte communities from 1,539 plant individuals, representing 401 vascular plant genera. To thoroughly quantify endophyte-plant specialization at each site, we sampled one individual of every co-occurring plant species within a 50 m² plot and sampled across five plots at each site (100 plots total).

**Figure 1.**
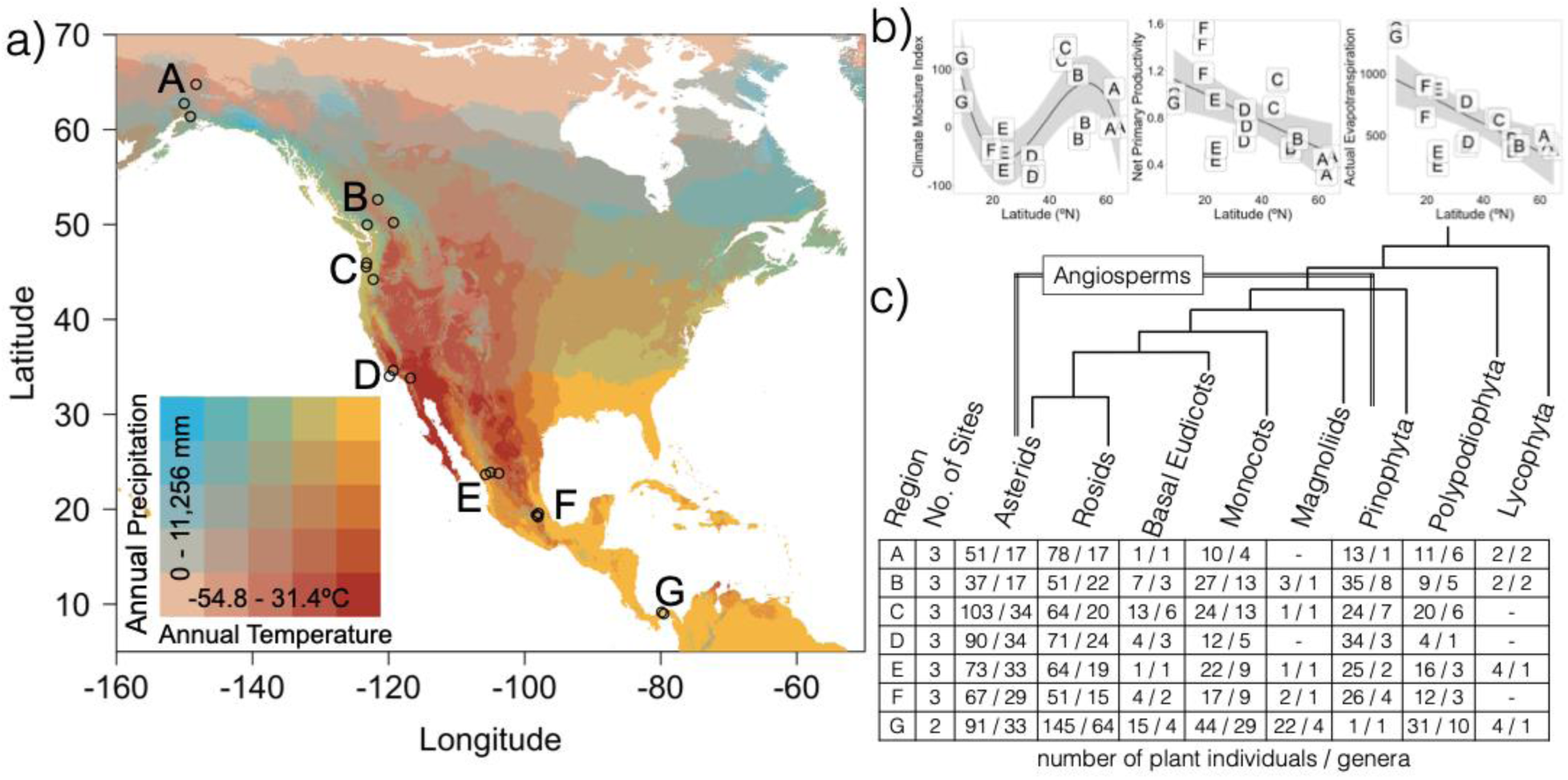
Foliar fungal endophyte communities were sampled across a 55° latitudinal gradient that spanned Central and North America. (a) Endophytes and plants were sampled from 20 sites that each contained five 50 m^2^ plots (n = 100; open circles) across seven sampling regions (A = Alaska, USA; B = British Columbia, Canada; C = Oregon, USA; D = California, USA; E = Durango, Mexico; F = Tlaxcala, Mexico; G = Panamá Oeste, Panama). Letters refer to sampling regions in panels b and c. (b) Climate moisture index (kg/m²/month) was high in the tropics and temperate rainforests. Net primary productivity (kg C/m²/year) and actual evapotranspiration (kg/m²/year) increased toward the tropics. (c) Table lists number of plant individuals (n = 1,539) and vascular plant genera (n = 401) in the final dataset. The cladogram of major plant lineages is based on Sessa et al. (2014) and Li et al. (2021).

In addition to network size, factors related to resource availability or access to host associations could influence specialization (Hurlbert 1978, Blüthgen et al. 2006). For example, differences in the physical and chemical properties of leaves among host species may act as barriers to the recruitment of symbiotic fungi, helping to explain why endophyte community composition often depends on the identities or the evolutionary relatedness of host plants (Liu et al. 2019, U’Ren et al. 2019, Whitaker et al. 2020, Darcy et al. 2020). To partially address these host trait differences, we included plant phylogenetic diversity in a model to predict specialization. Furthermore, because most fungal endophytes are passively dispersed through the environment, especially during periods of high moisture availability and when leaf stomata are open, the evolution and maintenance of their specialization may depend on the probability of colonizing the same host (Apigo and Oono 2022), leaf conditions that determine fungal spore viability (Darcy et al. 2020), and the local climate (Oita et al. 2021, U’Ren et al. 2024). Hence, we also included plant occurrence, net primary productivity (NPP), actual evapotranspiration (AEP), and a climate moisture index (CMI) in our model.

To test for LSG and its underlying drivers, we measured specialization of the endophyte-plant community in each plot by considering the presence and absence of associations (1-connectance) or the frequency and evenness of associations (H2’ specialization). Then, we examined how patterns of specialization varied across latitude after accounting for differences in network size (endophyte and plant richness), the phylogenetic diversity of plants, and plant occurrence across plots. Finally, we measured the degree to which specialization could be explained by endophyte richness, the plant community (plant richness, plant phylogenetic diversity, plant occurrence), AEP, NPP, CMI, or sampling date with a piecewise structural equation model (pSEM). Because our dataset includes all unique plant genera and endophytes per plot, leading to differences in network size across plots, we re-analyzed the pSEM using subsampled networks of equal size to control for its effect. Our study is the first to systematically and comprehensively, to the extent possible with amplicon sequencing, test for LSG in endophyte-plant communities and account for common ecological, evolutionary, and methodological factors known to shape measures of specialization.

## Materials and Methods

### Sampling design

Plant samples were collected from 20 sites that ranged in latitude from 8°N to 64°N, mean annual temperature from -1.55°C to 26.75°C, and mean annual precipitation from 422.6 kg m^-2^ year^-1^ to 3,305.2 kg m^-2^ year^-1^ during the summers of 2016 to 2017 (Figure 1; Table S1). Most sites were forests where the entire canopy was composed of conifers (n = 15; e.g., *Abies*, *Picea*, *Pinus*, *Pseudotsuga*, *Thuja*, *Tsuga*) or landscapes that contained conifer stands (n = 3; *Juniperus*, *Pinus*). The sites in Panama (n = 2) were broadleaf tropical rainforests mostly composed of angiosperms (e.g., *Fridericia*, *Machaerium*, *Piper*, *Psychotria*). Within each site, we haphazardly identified a general area that included dominant tree species and sampled from five 50 m^2^ plots using a random number table with directional headings (1-360°) and number of steps (1-100) to the first corner of a given plot. Within each plot, we haphazardly sampled from one individual of every plant species. We collected 10 leaves with a healthy appearance from plant individuals that were reachable by hand or with a pole clipper. For evergreen plants (e.g., conifers), we sampled leaves less than one year old to collect the cohort of fungi representative of that year’s growing season.

### Molecular and bioinformatic methods

Leaves were surface-sterilized by sequential immersion in 10% commercial bleach for 2 minutes, UV-treated deionized water for 30 seconds, and 70% ethanol for 2 minutes. Sterilized leaves were air-dried in a sterile laminar flow hood, frozen, and stored at -80°C until they were homogenized using a mortar and pestle with liquid nitrogen. DNA was extracted from 80 mg of plant tissue using a modified 2% CTAB method (Doyle and Doyle 1987) and purified with the DNA Clean and Concentrator-5 kit (Zymo Research). The internal transcribed spacer 1 (ITS 1) region was amplified from 10 ng of genomic DNA with fungal-specific primers ITS1F-KYO1 and ITS2-KYO1 (Toju et al. 2012) modified with Illumina overhang adapters (library preparation details can be found in Apigo and Oono 2022). The PCR products from each plant individual were sequenced on two separate Illumina MiSeq runs (n = 14 sequencing runs), using the same dual indices to evaluate potential biases associated with individual sequencing runs. Each run sequenced 250 base pairs of paired-end reads and included a 15% spike-in of PhiX (California NanoSystems Institute, University of California, Santa Barbara). Positive controls on each sequencing run included PCR4-TOPO TA vectors (Invitrogen) containing the ITS1 region from Ascomycota and Basidiomycota endophytes. These controls were used to identify reads affected by Illumina “index hopping,” where unexpected i5 and i7 primer combinations can occur (Huson et al. 2007, van der Valk et al. 2020).

Merged reads were trimmed of primers with Cutadapt v3.5 (Martin 2011). The ITS1 region was extracted with ITSx (Bengtsson-Palme et al. 2013) and reads were filtered and clustered into 97% Operational Taxonomic Units (OTUs) with de novo chimera identification using USEARCH v11.0.667 (Edgar and Flyvbjerg 2015) and VSEARCH v2.21.1 (Rognes et al. 2016). OTUs were queried against the GenBank database (Altschul et al. 1990) with BLAST+ v2.12.0 (Camacho et al. 2009). OTUs not identified within the kingdom Fungi were removed with MEGAN v6.22.2 (Huson et al. 2007). All bioinformatics were performed on the Center for Scientific Computing Knot cluster at the University of California, Santa Barbara. All statistical analyses were performed in R v4.4.2 (R Core Team 2022).

Endophyte communities that were amplified from the same genomic DNA but sequenced on different MiSeq runs were combined by summing read abundances. The vast majority of reads (mean across sequencing runs = 99.84% of reads) were correctly assigned to each positive control (n = 27; Table S5), suggesting that barcode switching and contaminations were minimal in this dataset. Endophyte communities were rarefied to a sequencing depth of 3,500 reads with the “rrarefy” function in vegan based on sample completeness curves (Figure S1) using the “iNEXT” function in the iNEXT (Interpolation and Extrapolation for Species Diversity) package (Hsieh et al. 2016).

The ribulose-1,5-bisphosphate carboxylase oxygenase (forward primer: rbcLa-F, reverse primer: rbcLajf634-R; Kress and Erickson 2007, Fazekas et al. 2008) or the internal transcribed spacer 2 (forward primer: ITS-u3; reverse primer: ITS-u4; Cheng et al. 2016) regions were sequenced with Sanger or Illumina MiSeq (250 paired-end reads) chemistry, respectively, and queried with BLAST to approximate plant identity (Figure 1; Table S2). Taxonomic ranks for each plant were standardized to NCBI taxonomy (ncbi.nlm.nih.gov/taxonomy).

Of 1,898 field-collected samples, we successfully amplified and sequenced plant and fungal DNA for 1,657 samples. Because currently available phylogenetic trees for plant genera in North and Central America only include vascular plants (Jin and Qian 2022), we removed non-vascular plants (118 plant samples) from all analyses in order to include plant phylogenetic distance or diversity as a covariate in statistical models. The final rarefied dataset comprised 5,383,870 reads across 18,732 non-singleton 97% OTUs and 1,539 plant individuals that comprised 401 vascular plant genera. Because we could not identify all plants to the species-level by observation and marker gene sequencing, plants that were identified to the same genus in a plot were combined into a single sample by summing the reads between them and were then rarefied. Combining the endophyte communities of plant congeners into a single sample versus randomly choosing one representative plant congener in each plot did not significantly change measures of community structure and specialization (Figure S2). After rarefying, endophyte OTUs with a relative read abundance <0.05% were excluded (OTUs with an absolute abundance of less than 24 reads, on average) in each plot. We chose this exclusion threshold based on the number of unexpected reads in the positive control samples.

### Measures of community structure

Endophyte alpha diversity was measured as the richness of 97% fungal OTUs within each plant sample with the “specnumber” function in the vegan package (Oksanen et al. 2020). Endophyte gamma diversity was measured as the number of unique endophyte OTUs across all plant samples in a plot. Within each plot, endophyte beta diversity based on Bray-Curtis dissimilarities was measured as the dispersion of endophyte communities among plant samples projected in principal coordinate space with the “betadisper” function in the vegan package. The turnover and nestedness components of endophyte beta diversity were separated with the “beta.pair” function in the betapart package (Baselga 2010).

Specialization was measured across the endophyte and plant community within each plot as 1-connectance with the “networklevel” function and the H2’ specialization index with the “H2fun” function in the bipartite package. Connectance measures the proportion of realized associations and H2’ measures the frequency and evenness of associations between endophyte OTUs and plant genera. We also measured other metrics of specialization that considered the specificity of endophyte OTUs to plant genera and vice versa: the presence-absence resource range index (Poisot et al. 2012) was measured with the “PDI” function and the abundance-weighted d’ specialization index (Blüthgen et al. 2006) was measured with the “dfun” function in the bipartite package. All specialization indices range from 0 (most generalized) to 1 (most specialized).

Because our plots varied in the number of endophyte and plant species (i.e., network size), which is known to bias metrics of specialization (Blüthgen et al. 2006), we standardized network size across plots. We subsampled a uniform number of plants (5 samples) and endophytes (15 OTUs) from each plot 1,000 times with replacement to create bootstrapped measures of specialization. We chose this network size (5 x 15) based on the minimum number of plant samples and the maximum number of endophyte OTUs that could be extracted from a random subsampling event. To control for differences in read counts, we divided read counts by the sum of read counts in each subsampled network such that each network summed to one. We constructed statistical models with 1-connectance, H2’, bootstrapped 1-connectance, or bootstrapped values of H2’ as the response variable to assess whether differences in network structure biased our inferences.

Plant alpha diversity was measured as the number of unique plant genera in a plot. Plant gamma diversity was measured as the number of unique plant genera in a site. Plant beta diversity was measured as the dispersion of plots relative to their site centroid in principal coordinate space using the Jaccard index.

### Latitudinal patterns of community structure

Measures of community structure (endophyte alpha diversity, endophyte beta diversity, endophyte gamma diversity, endophyte-plant specialization; Figure 2a) were regressed as a function of latitude with linear or polynomial mixed-effects models with the “lme” function in the nlme package (Pinheiro and Bates 2022). We selected the best-fit model with AICc, refit the chosen model with restricted maximum likelihood, and measured marginal R-squared coefficients (Nakagawa and Schielzeth 2013) with the “model.sel” and “r.squaredGLMM” functions in the MuMIn package (Barton 2009). For each measure of community structure, we included endophyte and plant-related covariates and re-tested the significance of the latitude term(s) using marginal sums of squares (Table S3). We tested for spatial autocorrelation using Moran’s I with the “testSpatialAutocorrelation” function in the DHARMa package (Hartig 2020). In a separate analysis, we performed linear matrix regression (999 permutations) with the “mantel” function in the vegan package (Oksanen et al. 2020) to understand how the effect of plant phylogenetic distance on endophyte composition varied across latitude.

**Figure 2.**
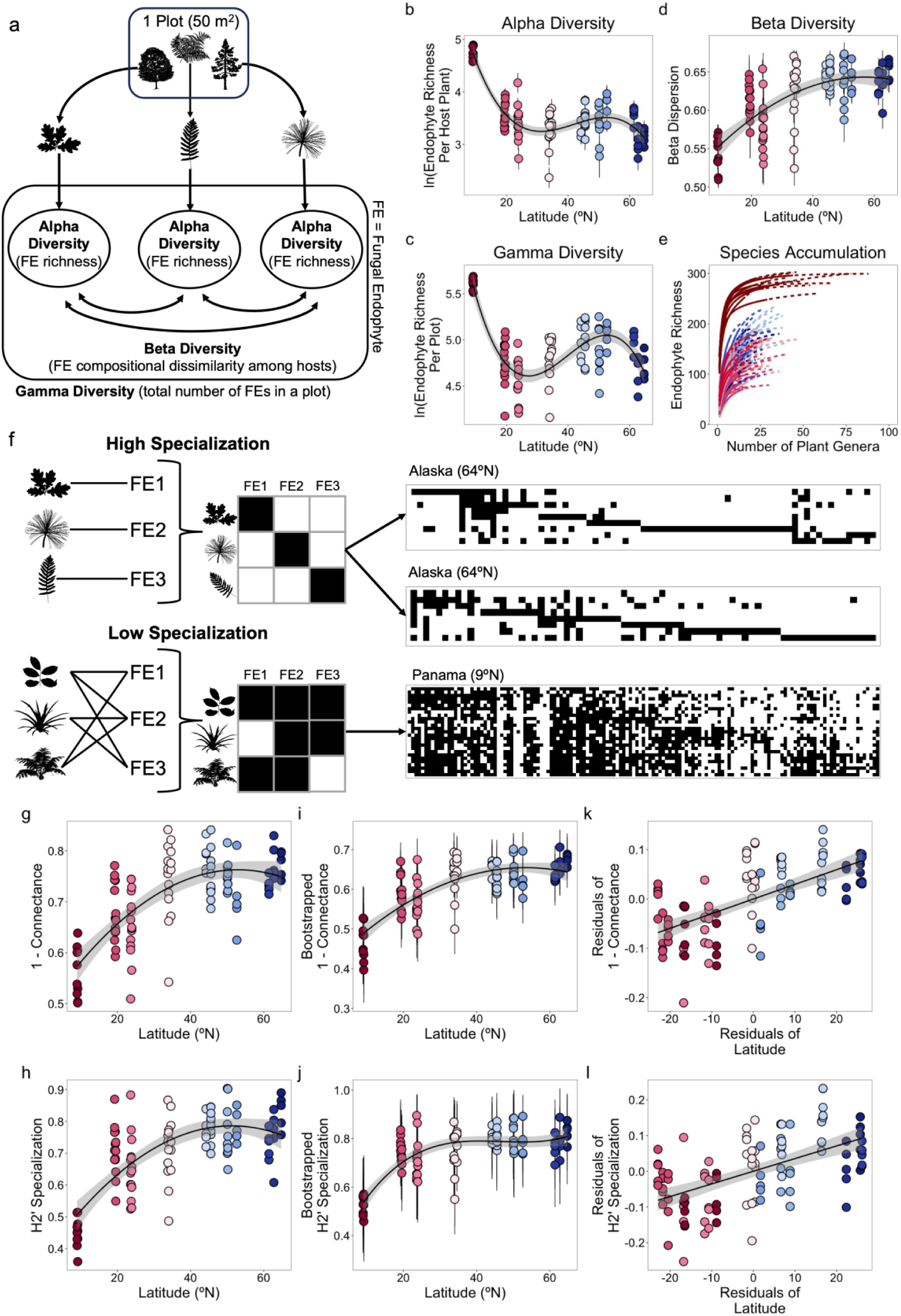
Foliar fungal endophytes are most species-rich and most generalized in their associations with plants in tropical forests. (a) Conceptual diagram of endophyte alpha, beta, and gamma diversity at a plot scale (n = 100). (b) The richness of 97% endophyte OTUs per plant sample (alpha diversity) is greatest in the tropics (R^2^ = 0.69; cubic latitude term: p < 0.001). (c) The total number of endophytes per plot (gamma diversity) is greatest in the tropics (R^2^ = 0.61; cubic latitude term: p < 0.001). (d) Endophyte communities are most compositionally similar among plant samples (Bray-Curtis dissimilarity) in the tropics (R^2^ = 0.49; quadratic latitude term: p = 0.019). (e) Tropical rainforests in Panama (dark red lines) harbor a greater richness of endophytes than temperate and subtropical coniferous forests (blue and pink lines) at any sampling depth of the plant community. Each species accumulation curve represents one plot. Solid lines are interpolated values and dotted lines are extrapolated values. (f) Conceptual diagram (left) and representative examples of communities with high or low specialization (right). Black squares represent the presence of an endophyte-plant association, while white spaces represent the absence of an association. (g) The proportion of realized associations between endophytes and plants (1-connectance) increases toward the tropics (R^2^ = 0.51; quadratic latitude term: p = 0.0068). (h) Endophytes and plants form more numerous and more symmetric associations (H2’) toward the tropics (R^2^ = 0.50; quadratic latitude term: p = 0.0012). (i, j) Bootstrapped values of specialization standardized by the number of endophyte OTUs, the number plant samples, and the sum of read counts in each subsampled network (n = 1000 subsamples per plot) support the same trends as non-bootstrapped values of specialization. (k, l) Partial regression plots show the decline in specialization toward the tropics after accounting for within and between site effects of endophyte richness, plant richness, plant phylogenetic diversity, and plant occurrence. Each point represents the mean or single value of community structure per plot (n = 100) and colors refer to latitude. Greater values correspond to greater endophyte diversity or specialization. Bars refer to standard error in panels b, c, and d and bars refer to standard deviation in panels i and j. Images of plants in panels a and f are from phylopic.org.

### Structural equation modeling

We used a piecewise structural equation model (pSEM) to test the direct effects of endophyte richness, plant community structure (plant richness, plant phylogenetic diversity, and plant occurrence), landscape vegetation factors (actual evapotranspiration and net primary productivity), climate moisture index, and sampling date (day and year) on specialization (Figure 3). The pSEM considered the indirect effects of plant community structure, landscape vegetation factors, climate moisture index, and sampling date on specialization through their effects on endophyte richness. The pSEM also considered the indirect effects of sampling date and climate moisture index on specialization through their effects on plant community structure, and the indirect effect of climate moisture index on specialization through its effect on landscape vegetation factors. We considered the effect of sampling date on endophyte richness, plant community structure, and specialization because these values were dependent on samples collected in the field. Evapotranspiration and productivity were not dependent on sampling date because these values are averages of yearly values from an online database. Thus, sampling date was not included as a path to evapotranspiration and net productivity. In pSEM, each relationship between a response variable and its associated predictor variable(s) was estimated as a separate linear regression or a linear mixed effects model.

**Figure 3.**
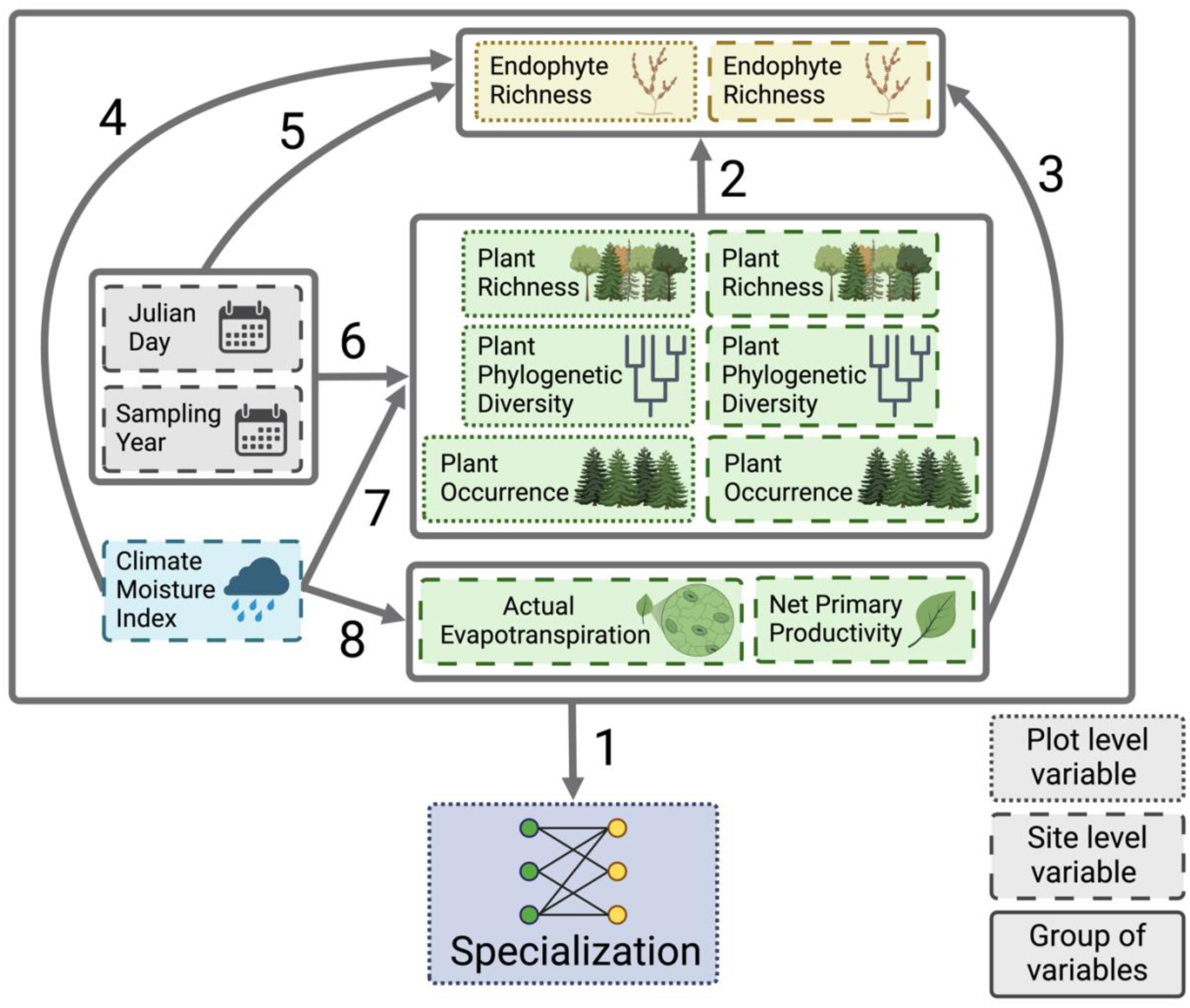
Diagram of hypothesized relationships that determine the latitudinal specialization gradient for foliar fungal endophytes and plants. The structural equation model tested the direct effects of endophyte richness, plant community structure (plant richness, plant phylogenetic diversity, and plant occurrence), landscape vegetation factors (actual evapotranspiration and net primary productivity), climate moisture index, and sampling date (day and year) on specialization (arrow 1). The model considered the indirect effects of plant community structure (arrow 2), landscape vegetation factors (arrow 3), climate (arrow 4), and sampling date (arrow 5) on specialization through their effects on endophyte richness. The model also considered the indirect effects of sampling date (arrow 6) and climate moisture index (arrow 7) on specialization through their effects on plant community structure, and the indirect effect of climate moisture index on specialization through its effect on landscape vegetation factors (arrow 8). We considered the effect of sampling date on endophyte richness (arrow 5), plant community structure (arrow 6), and specialization (arrow 1) because these values were dependent on samples collected in the field. Evapotranspiration and productivity were not dependent on sampling date because these values are averages of yearly values from an online database. Solid grey line boxes are groups of predictor variables and arrows connecting solid grey line boxes to other variables represent all possible relationships between these groups. Dashed line boxes are site level predictor variables that were measured at the site level or are site level means (i.e., the between-site effects; see Methods), while small dotted line boxes are plot level predictor variables standardized to site level means (i.e., the within-site effects; see Methods) or the plot level response variable (specialization; H2’ index). The model also considered correlated errors among the six plant community structure and the two landscape vegetation variables and a correlation between the two endophyte richness variables, but arrows among these boxes are not shown for visual simplicity.

Sampling day and year were included as covariates to account for intra-annual variation in sampling across 115 days (172-287 Julian days) and inter-annual variation in sampling across two years (2016 and 2017). Climate moisture index (kg/m^2^/month), a measure of water availability (mean annual precipitation) relative to water demand (potential evapotranspiration), was extracted from Climatologies at high resolution for the Earth’s land surface areas (version 2.1; Karger et al. 2017, 2020). Endophyte richness was measured as the number of unique endophyte OTUs across all plant samples in a plot. Plant richness was measured as the number of vascular plant genera within a plot. Endophyte and plant richness were natural log scaled. We used V.Phylomaker2 to extract phylogenetic distances for vascular plant genera (Jin and Qian 2022) and calculated the standardized effect size of the mean pairwise phylogenetic distance per plot with the “ses.mpd” function in the picante package (Kembel et al. 2010). Plant occurrence was measured as the frequency (1 to 5 plots) of each plant genus across the five plots at a given site and averaged at the plot level. Plots composed of plants that were more frequent within a site had lower compositional differences relative to their site’s centroid and thus lower plant beta diversity (Pearson’s r: -0.64, p < 0.001; Figure S3d). We chose to use plant occurrence over plant beta diversity because we aimed to test whether host commonness, and thus host availability, was associated with greater specialization. Actual evapotranspiration (kg/m²/year) and net primary productivity (kg C/m²/year) were measured by the Moderate Resolution Imaging Spectroradiometer (modis.gsfc.nasa.gov) at a 500-meter resolution and extracted using AρρEEARS v6.1 (appeears.earthdatacloud.nasa.gov).

When predictor variables that were measured at the plot level (endophyte richness, plant richness, plant phylogenetic diversity, plant occurrence) were included in a linear regression, we treated site as a random effect because five plots were nested within each site. We accounted for within-site and between-site effects of these predictor variables on specialization or endophyte richness (Group Mean Centering Design; Byrnes and Dee 2025). Between-site effects were calculated as the mean of the five plot level measures at each site and the within-site effects were calculated by subtracting the site level mean from each plot level measure. Variables were then scaled using the “scale” function. We used the H2’ specialization index as the response variable. We repeated the analysis with bootstrapped values of H2’ that controlled for the number of endophyte OTUs, the number of plant genera, and the sum of read counts in each subsampled network. The H2’ index was not spatially autocorrelated (Moran’s I: p > 0.05; Table S3) and thus, we did not include a spatial correlation structure in the pSEM.

We fit an initial model with the “psem” function in the piecewiseSEM package (Lefcheck 2016), then used backwards elimination to avoid overfitting the model (Pinheiro et al. 2023). At each step, we dropped the path with the highest p-value, compared nested pSEMs with AICc, and repeated this procedure until the AICc of the simpler pSEM was within two delta AICc of the more complex pSEM (Table S4). We modeled relationships among plant factors (e.g., evapotranspiration and net primary productivity) as correlated errors where no directionality was assumed and tested their independence from the pSEM with the “dSep” function in the piecewiseSEM package (Table S4). We used Fisher’s C as goodness-of-fit test to evaluate whether the hypothesized structure of the pSEM adequately explained the observed data. We report standardized regression coefficients (β) that correspond to the strength of each predictor, relative to other covariates, in the pSEM. We constructed partial regression plots to visualize the strength of the relationship between a predictor variable and specialization while accounting for other covariates in the model (Moya-Laraño and Corcobado 2008).

## Results

### The tropics have the greatest endophyte alpha diversity and the lowest endophyte beta diversity

Endophyte richness displayed a non-linear pattern with latitude, with endophyte alpha diversity (per plant sample; Figure 2a, 2b; Table S3a) and endophyte gamma diversity (per plot; Figure 2a, 2c; Table S3b) peaking in the tropical rainforests of Panama (8-10°N). Moving northward, endophyte richness declined from Panama to Mexico (19-23°N), followed by a slight rebound from southern California (33-35°N) to Oregon and British Columbia (35-50°N), before decreasing again at the most northern site of Alaska (61-65°N). These trends reveal a cubic relationship between endophyte richness and latitude (Figure 2b-c), highlighting tropical rainforests as a diversity hotspot within this gradient. Endophyte richness was consistently highest in the tropics regardless of number of hosts sampled per plot (Figure 2e) or the plant lineage (Figure S4).

Endophyte community composition among host samples (beta diversity; Figure 2a) was most similar in the tropics and most different at higher latitudes (Figure 2d). This pattern was driven primarily by reduced species replacement (i.e., turnover, Figure S5a) rather than by lower species loss (i.e., nestedness, Figure S5b) in the tropics. After accounting for covariates, such as endophyte richness, plant richness, plant phylogenetic diversity, and plant occurrence, endophyte beta diversity remained lowest in the tropics (Table S3c).

### Endophyte-plant specialization decreases toward the tropics

Specialization between endophytes and plants, measured as either the presence and absence of associations (1-connectance) or evenness (H2’), decreased toward the tropics (Figure 2g-h). These metrics were correlated (Pearson’s r: 0.73, p < 0.001), indicating that communities with more realized associations had species with more overlapping interactions. These patterns held even after subsampling communities to uniform network sizes and standardized read counts (Figure 2i-j) and when accounting for endophyte and plant community structure, such as endophyte richness, plant richness, plant phylogenetic diversity, and plant occurrence (Figures 2k-l, Table S3d-e). A key driver of this trend appears to be lower host specificity (i.e., resource range index and d’) among endophytes in the tropics: on average, each endophyte associated with a greater proportion of the plant community (Figure S6a-b). For example, the average endophyte OTU was found in 43.1% of host samples in a Panamanian plot (9.11 plant samples on average), compared to 22.6% of host samples (2.06 plant samples on average) in an Alaskan plot (Figure 2f). Similarly, plant hosts of Panama formed more generalized associations with endophytes, associating with a larger proportion of the endophyte community (Figure S6c-d). Endophyte beta diversity was positively correlated with 1-connectance (Pearson’s r: 0.76, p < 0.001) and H2’ specialization (Pearson’s r: 0.92, p < 0.001).

The latitudinal decline in specialization toward the tropics could not be attributed to host phylogenetic distance, as the relationship between plant phylogenetic distance and endophyte dissimilarity within each plot did not decrease toward the tropics (Figure S5d). Additionally, while communities with rare endophytes (i.e., low read abundances) might seem more specialized due to their limited occurrence in host samples, plots with a greater number of rare endophytes were not more specialized than those with abundant ones (Figure 6). Hence, read abundance did not influence measures of specialization. Changes in endophyte-plant specialization were not strongly linked to heterogeneity of plant communities, as plant beta diversity among plots per site, which correlated with plant occurrence (Pearson’s r: -0.64, p < 0.001; Figure S3d), did not significantly vary across latitude (Figure S3c). Shifts in the relative abundance of fungal taxonomic class coincided with the decline in specialization toward the tropics: Sordariomycete fungi increased in relative abundance, while Leotiomycete fungi decreased in relative abundance toward southern latitudes (Figure S7).

### Climate and plant community features, not endophyte richness, drive specialization between endophyte and plant communities

Actual evapotranspiration (AEP) emerged as the strongest driver of specialization, followed by plant richness, climate moisture index (CMI), net primary productivity (NPP), and sampling day (Figures 3-5; Table S4). Communities with higher AEP and greater plant richness were less specialized (paths a and b in Figure 4, Figure 5a-b). For example, the two sites in Panama, which exhibited the lowest values of specialization (Figure 2h), also had the highest plant richness (Figure S8h) and AEP (Figure S8l). In contrast, communities in areas with higher CMI (i.e., greater precipitation than evaporative demand) and higher NPP (i.e., greater plant biomass) were more specialized (paths c and d in Figure 4, Figure 5c-d). This trend was evident in temperate rainforests of Oregon, characterized by high CMI (Figure S8c), and coniferous forests in Mexico, characterized by high NPP (Figure S8m), which had more specialized communities. Sampling day also influenced specialization; sites sampled later in the growing season had lower specialization (path e in Figure 4, Figure 5e, Figure S8a).

**Figure 4.**
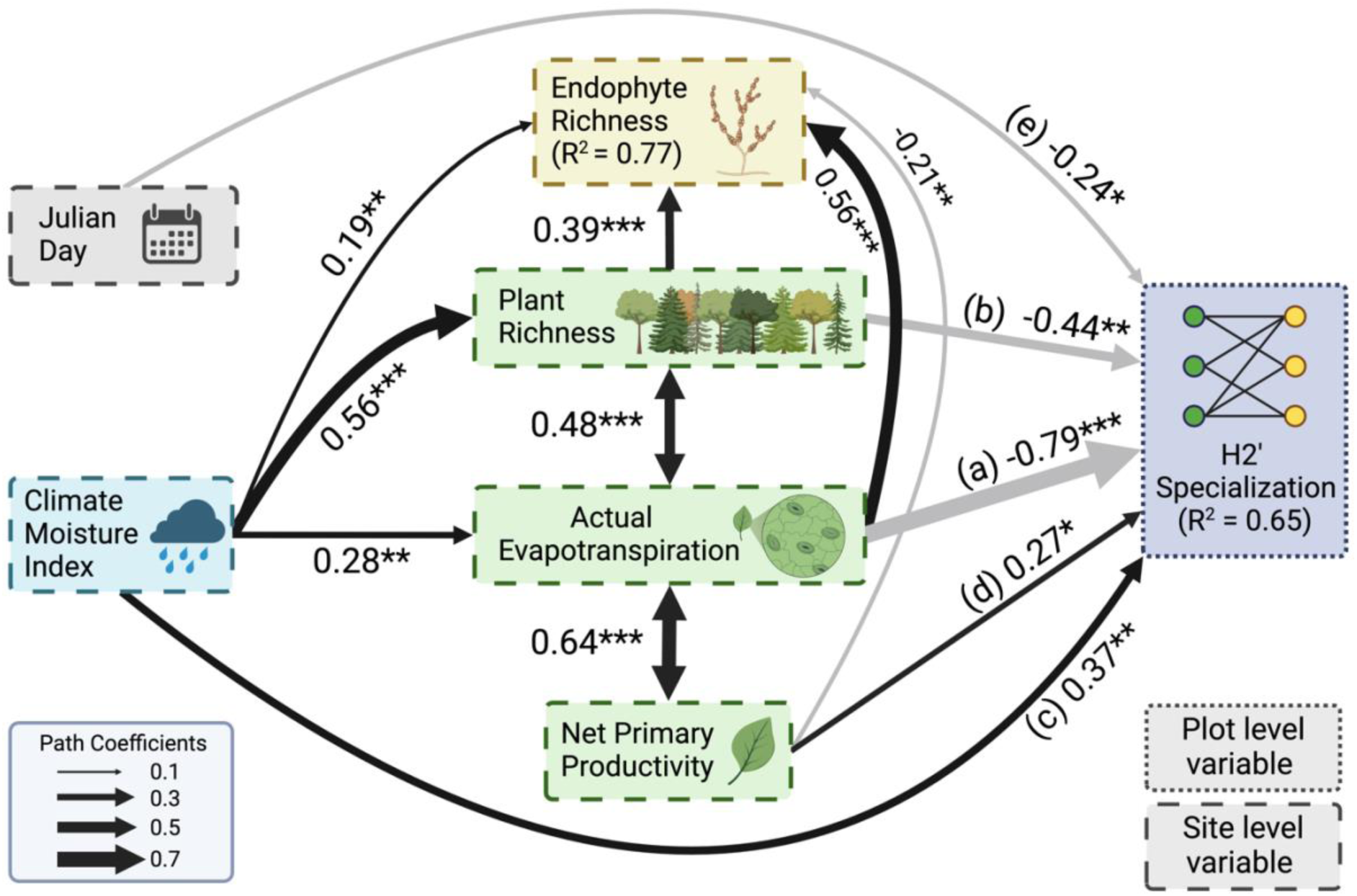
Climate and plant community features, not endophyte richness, drive the latitudinal specialization gradient for endophytes and plants. Black and grey paths distinguish positive and negative relationships, respectively, in the structural equation model (Fisher’s C = 58.78, df = 78, p = 0.95). Unidirectional paths represent relationships in a simple or multiple linear regression, while bidirectional arrows represent correlated errors between variables. The width of each path corresponds to the magnitude of the standardized regression coefficient or the correlation coefficient. Paths shown are labeled with their statistical significance (***: 0 < p < 0.001, **: 0.001 < p < 0.01, *: 0.01 < p < 0.05). R-squared values are the marginal r-squared values from linear mixed effect models. Boxes with dotted or dashed lines are plot or site level variables, respectively (see Figure 3). Paths between predictor variables and specialization that were not statistically significant are not shown. Full model results can be found in Table S4. These relationships held with a model using a bootstrapped H2’ index (Figure 2j) that controlled for differences in network size (Figure S9).

**Figure 5.**
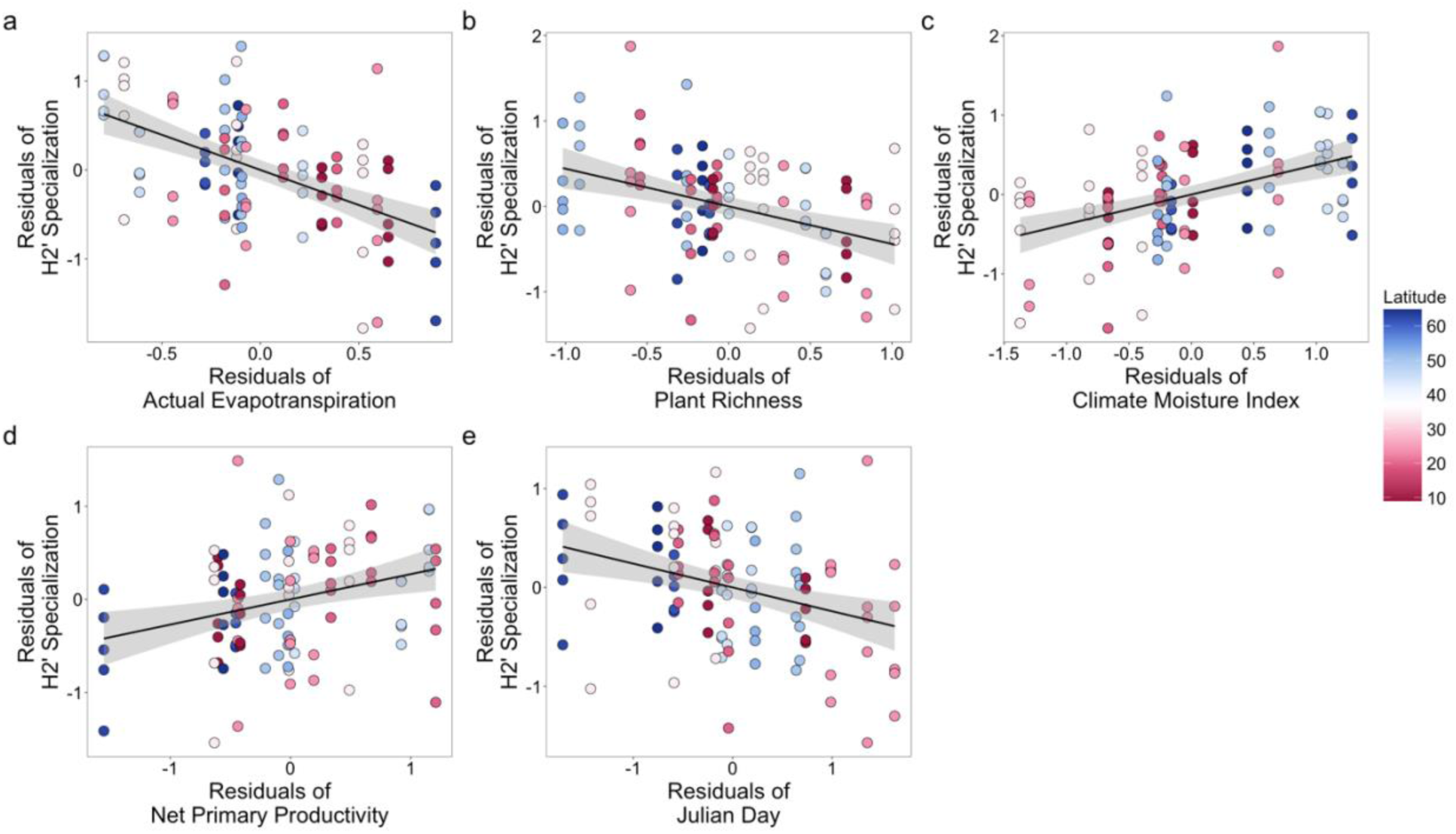
Relationships between H2’ specialization and predictors in the structural equation model. Each panel is a partial regression plot that shows how H2’ specialization varies with respect to a predictor variable while controlling for other covariates in the structural equation model (Figure 4; Table S4). Each point is one plot (n = 100) and colors refer to latitudinal position. (a) Communities with greater evapotranspiration were less specialized (β = - 0.79, p = 0.0003). (b) Communities with greater plant richness were less specialized (β = -0.44, p = 0.0049). (c) Communities with greater water availability were more specialized (β = 0.37, p = 0.0014). (d) Communities with greater primary productivity were more specialized (β = 0.27, p = 0.023). (e) Sites sampled later in the year were less specialized (β = -0.24, p = 0.013).

**Figure 6.**
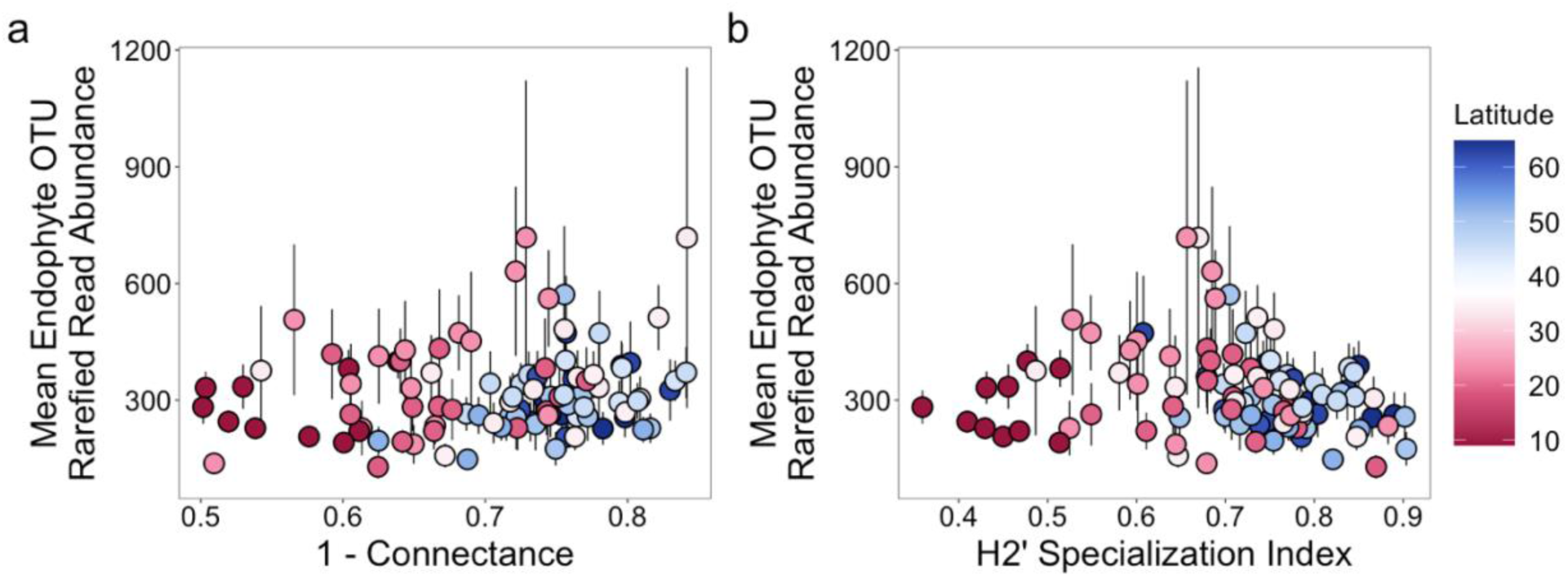
Plots composed of rarer endophyte OTUs were not biased to have greater specialization than plots with more abundant endophyte OTUs. (a, b) Rarefied endophyte OTU read abundances averaged at the plot level were not negatively correlated with 1-connectance (Pearson’s r: 0.20, p = 0.046) or the H2’ specialization index (Pearson’s r: -0.15, p = 0.13). Each point is a plot, bars refer to standard error, and colors refer to latitudinal position.

Importantly, the piecewise structural equation model (pSEM) with H2’ values from subsampled networks of uniform sizes (see Methods; Figure S9) produced regression coefficients similar to those in the non-boostrapped H2’ model (Figure 4), demonstrating the robustness of these results to differences in network scaling. The pSEM was also robust to the exclusion of the two sites in Panama (Figure S10) and thus these results are not solely dependent upon the high AEP, high plant richness, and high endophyte richness found in tropical rainforests. The pSEM was designed to assess how plot-level predictors, including endophyte richness, plant richness, plant phylogenetic diversity, and plant occurrence, influenced specialization within and between sites (Figure 3). However, only between-site effects on specialization were statistically significant in the final pSEM model (Figure 4; Table S4), underscoring the role of broader environmental differences in shaping specialization patterns.

## Discussion

We found that fungal endophyte species richness peaks in the tropics as expected by the latitudinal diversity gradient, following trends of their host plants (Figure 2b-c; Figure S3). However, endophyte-plant specialization did not align with the latitudinal niche-breadth hypothesis (Figure 2g-l; MacArthur 1972, Schemske et al. 2009), which is often invoked to explain the high species richness in the tropics (Dobzhansky 1950, MacArthur 1965, Janzen 1970, Connell 1971, Dyer and Forister 2019). We found that endophytes and plants formed a greater number of more symmetric associations toward the tropics and therefore were the most generalized in areas with the greatest richness of endophytes and plants (Figure 2g-l). The latitudinal specialization gradient has faced growing scrutiny (Novotny et al. 2002, Vázquez and Stevens 2004, Schleuning et al. 2012, Moles and Ollerton 2016), with increasing evidence that high specialization in the tropics is more the exception than the rule.

While the tropics contained the highest plant richness and highest endophyte richness, this diversity cannot be explained by narrower niches and greater specialization. Lower specialization was associated with greater plant richness (Figures 4-5), particularly in plots where host species occurred at lower frequencies within a site (Table S4). This negative association may be explained by the fact that most endophytes are horizontally transmitted among hosts (Rodriguez et al. 2009), and thus rely on the accessibility of compatible hosts to evolve host-specific symbioses. In species-rich tropical landscapes, endophyte propagules may be less likely to encounter the same host species across space and time. Hence, similar to how resource fragmentation limits specialist persistence (Janzen and Pond 1975, Kindlmann et al. 2007), a diverse plant community may functionally fragment host availability for specialist endophytes, promoting generalists in such landscapes. Although hosts with lower occurrence were associated with lower specialization indirectly through plant richness, supporting our explanation, we did not find a significant relationship between host occurrence and specialization in our model (Table S4). This could be because our measurement of host occurrence metric, based on 50 m² plots, does not adequately capture the spatial continuity of host distributions or dispersal scales relevant to shaping host-specific associations.

In addition to spatial constraints on host availability, host traits, such as leaf chemistry or morphology, which can correlate with evolutionary relatedness of hosts, may also influence patterns of plant-endophyte specialization across latitudes. Host identity and leaf traits have been shown to correlate with endophyte community composition (Christian et al. 2020) and can explain why closely related hosts tend to harbor more similar symbionts (Liu et al. 2019). Host identity and host phylogenetic distance were also more strongly associated with differences in endophyte composition than geography or other environmental covariates at both high and low latitudes (U’Ren et al. 2019, Darcy et al. 2020). However, in our study, neither plant phylogenetic dissimilarity nor diversity varied with latitude or influenced specialization (Figure S5d, Table S4). While host identity or host phylogenetic distance can explain why some endophyte-plant communities are more specialized than others, they did not explain the increase in specialization toward temperate latitudes.

The effects of endophyte dispersal and host trait filtering on specialization may also be influenced by the leaf environment. We found that higher actual evapotranspiration (AET), which reflects higher water loss through stomata—surface pores on leaves through which fungal endophytes are hypothesized to enter—was strongly associated with lower specialization (Figures 4-5). Because fungal endophytes can enter leaves through open stomata (Huang et al. 2018), prolonged stomatal opening or greater stomatal area (Wang et al. 2015, Liu et al. 2024) influenced or selected by high evapotranspiration, like in the tropical rainforests (but see Liu et al. 2018), may reduce host filtering and facilitate the recruitment of a greater richness of fungal species (Figure 2b-c, Figure 4). This could be especially pronounced in environments where airborne fungal spores are abundant, like in the understory of Panamanian rainforests (Arnold and Herre 2003, Arnold and Engelbrecht 2007). More frequent stomatal opening under greater AET might lead to more indiscriminate leaf colonization by airborne fungal spores, leading to higher endophyte-plant generalism (Figure 2g-l). Our findings align with previous work that has linked water loss and stomatal function to endophyte abundance and composition (Arnold and Engelbrecht 2007, Darcy et al. 2020), but further work that isolates the effect of stomatal physiology from climatic variation on patterns of specialization is necessary.

Like AET, climate moisture index (CMI) was positively associated with greater endophyte richness (Figures 4-5). This pattern has been observed in other studies (U’Ren et al. 2019, Oita et al. 2021) and likely reflects the role of water availability in fungal fitness via development of fruiting bodies, spore germination, and hyphal growth. Unlike AET, greater CMI was associated with greater specialization. Since the pSEM considers the indirect effect of CMI on specialization through AET (Figures 3-4), this result indicates that greater water availability independent of water loss promotes greater specialization. Interestingly, CMI is negatively correlated with precipitation seasonality (Pearson’s r: -0.50, p < 0.001), suggesting that more stable moisture regimes may allow for the evolution of specialization. Together, our results suggest that AET and CMI increase endophyte richness through different routes: greater AET may increase the richness of generalist endophytes by increasing colonization opportunities, while greater CMI may increase the richness of specialist endophytes by supporting the production and transmission of host-specific inoculum in consistently moist environments.

Like CMI, greater net primary productivity (NPP) was associated with greater specialization (Figures 4-5), which aligns with expectations of the latitudinal niche-breadth hypothesis that greater resources (i.e., plant biomass) could lead to greater specialization (MacArthur 1972, Vázquez and Stevens 2004). However, greater NPP was associated with lower endophyte richness (Figure 4). Although more productive environments are generally expected to support higher species richness, high resource settings may instead reduce richness by promoting fast-growing species that competitively exclude others (Grime 1973, Mittelbach et al. 2003, Brun et al. 2019). If competitively dominant endophytes are driving the negative relationship between NPP and endophyte richness, endophytes with narrower host ranges may have greater competitive ability on the few host species they associate with, which aligns with greater specialization in these ecosystems (Figures 4-5). However, the effects of NPP on both endophyte richness and specialization were relatively modest compared to other predictors in the model (e.g., AET). This may have occurred because the localized nature of most endophyte infections (Herre et al. 2007) allows competitively dominant species to coexist alongside others within the same leaf even in highly productive leaf environments.

Lastly, we found that communities sampled earlier in the growing season were more specialized than those sampled later in the growing season (Figures 4-5). One explanation for this relationship is a spurious correlation where some sites with lower specialization were sampled later, while some sites with higher specialization were sampled earlier (Figure S8a). However, endophyte communities tend to be more variable in composition and less specialized at earlier stages of leaf phenology (VanWallendael et al. 2022). Another explanation is that changes that occur over the lifespan of a leaf influence specialization or that later sampling dates excluded early season, short-lived plants that may have already senesced, skewing the sampled plant community. Nevertheless, whether greater variability in fungal composition earlier in the season or among short-lived plants is a general pattern is still an open question. Thus, the modest negative relationship between sampling date and specialization may reflect the order in which sites were sampled as well as biotic and abiotic features that covary with sampling date and were not captured by our model. Another concern is that the relationships identified by the structural equation model are dependent on the two Panama sites that had high plant richness, high AET, high NPP, and high CMI, which also had high endophyte-plant generalism. However, after excluding Panama, these relationships held (Figure S10), indicating that drivers of specialization in this study are not entirely driven by later sampling and environmental conditions unique to tropical rainforests.

The expectation that species associations are most specialized in the tropics has informed ecological theory in profound ways. For example, pathogens specialized to tropical plants have long been thought to promote tree diversity in the tropics by causing increased mortality among nearby conspecific trees (Janzen 1970, Connell 1971). Yet, like many paradigms in ecology, ecologists are gaining a greater understanding that these conceptual models are more complex than initially thought (e.g., Hülsmann et al. 2024). While we found that specialization decreased toward the tropics, contrary to expectations, we also find that specialization is not necessary for high species coexistence in the tropics: many species of endophytes coexist despite sharing many of the same host plants. Furthermore, elements of the latitudinal niche-breadth hypothesis, that more stable or productive environments can lead to some specialization, may still be applicable in explaining the global distribution plant-endophyte specialization. Our model provides a future framework to explore how the leaf environment as well as endophyte dispersal and host filtering could shape specialization. Further work that incorporates how fungi contend with the physical and chemical leaf landscape among host species (González-Teuber et al. 2021) and whether specialization depends on the functional roles of endophytes (Schulz and Boyle 2005) will be crucial to understanding the mechanisms that underlie how specialization is shaped in plant-endophyte communities.

## Data availability

Data and code associated with this study are archived in Figshare (https://figshare.com/s/14719ec6a9e4905b6f55). Sequencing data are archived in the National Center for Biotechnology Information Sequence Read Archive database (BioProject number PRJNA356423).

## Acknowledgements

This study and AA were supported by National Science Foundation Award 1841715, the University of California Santa Barbara Olivia Long Converse Fellowship, the University of California President’s Dissertation Year Fellowship, the Mycological Society of America Graduate Fellowship, and the Short Term Smithsonian Tropical Research Fellowship. RBPP is funded by São Paulo Research Foundation (FAPESP, 2025/01707-8 and 2025/29154-2). LCM acknowledges support from SNI SENACYT Program. We acknowledge Edward Allen Herre for field work and feedback on the manuscript. We acknowledge the following field sites and governing bodies for access to field sites: Denali State Park, Eklutna State Park, Bonanza Creek LTER, Garibaldi Provincial Park, Kalamalka Provincial Park, Quesnel River Research Center, HJ Andrews LTER, Clatsop State Park, Tillamook State Park, University of California Santa Cruz Island Reserve, University of California James San Jacinto Reserve, Los Padres National Forest, Secretaría de Medio Ambiente y Recursos Naturales México, and the Smithsonian Tropical Research Institute. We thank Cindy Quach, Kaylene Truong, Kiana Lee, Vanessa Greenman, Kevin Feller, Theodore Kwan, Megan Jung, Meggie Dinh, Carina Salaiz, Derek Tang, Simer Chima, Angelina Hyunh, Helen Chang, Thomas Masterson, Gabriela Dutra-Clarke, Hannah Hirou, Abby Camenisch, Alecsa Burwell, Jacob Harwood, Stephanie Austin, Gina Dempster, Helen Chang, Tiffany Wong, Serena Thai, Emily Lutz, Tessa Chou, Jules Goldstein, Abhishek Dighe, Angela Chu, Dulce Simental, Gokul Deep, Lindsey Calva, Jade Ingram, and Derek Fitzgerald for their efforts in the field and lab. The authors acknowledge the use of the Biological Nanostructures Laboratory within the California NanoSystems Institute, supported by the University of California, Santa Barbara and the University of California, Office of the President. We acknowledge computing support from the Center for Scientific Computing from the CNSI, MRL: an NSF MRSEC (DMR-1720256) and NSF CNS-1725797.

## Supplementary figures

**Figure S1.**
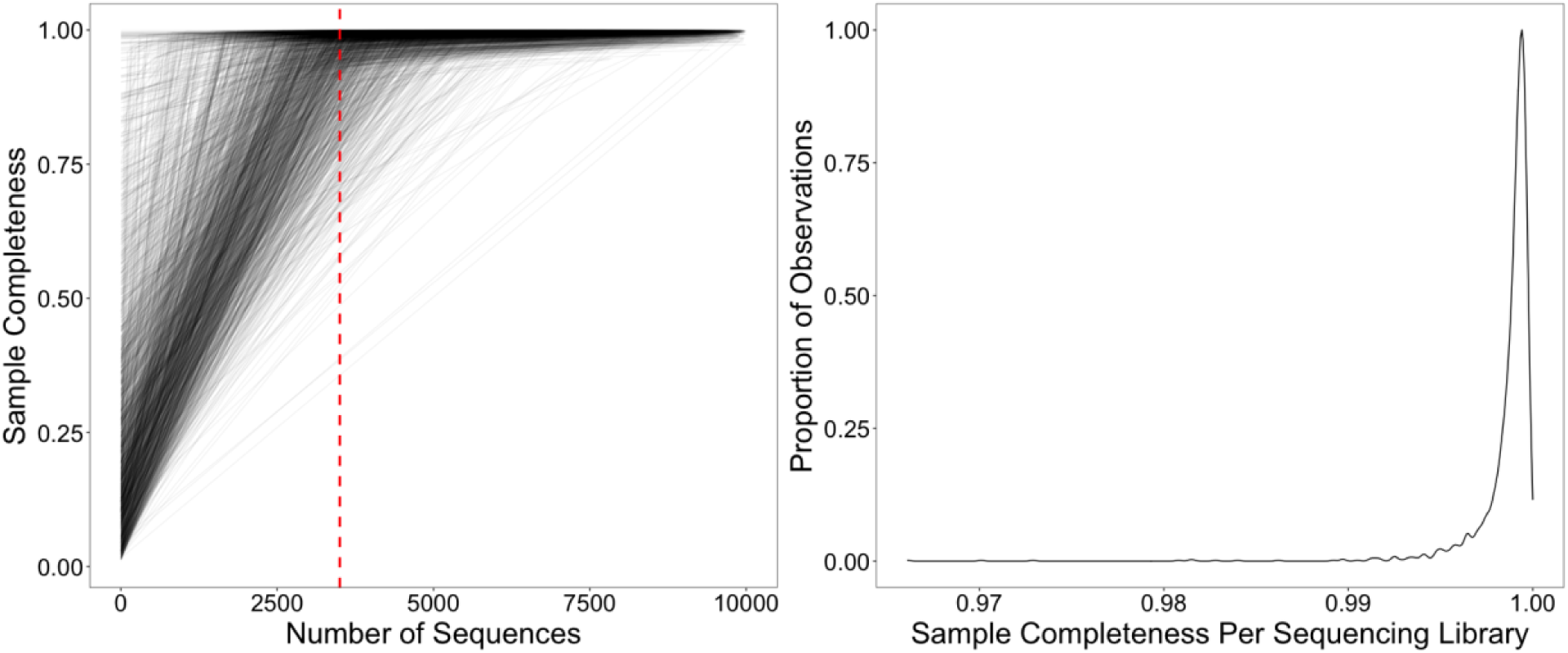
Sample completeness, a metric for how well an endophyte community was characterized by sequencing, in relation to (panel a) sequencing depth or as a (panel b) histogram. (a) Endophyte communities (each black line) were rarefied to a sequencing depth of 3500 reads (vertical red dashed line) because sample completeness tended to asymptote near that threshold. (b) The majority of the endophyte communities had sample completeness >99%.

**Figure S2.**
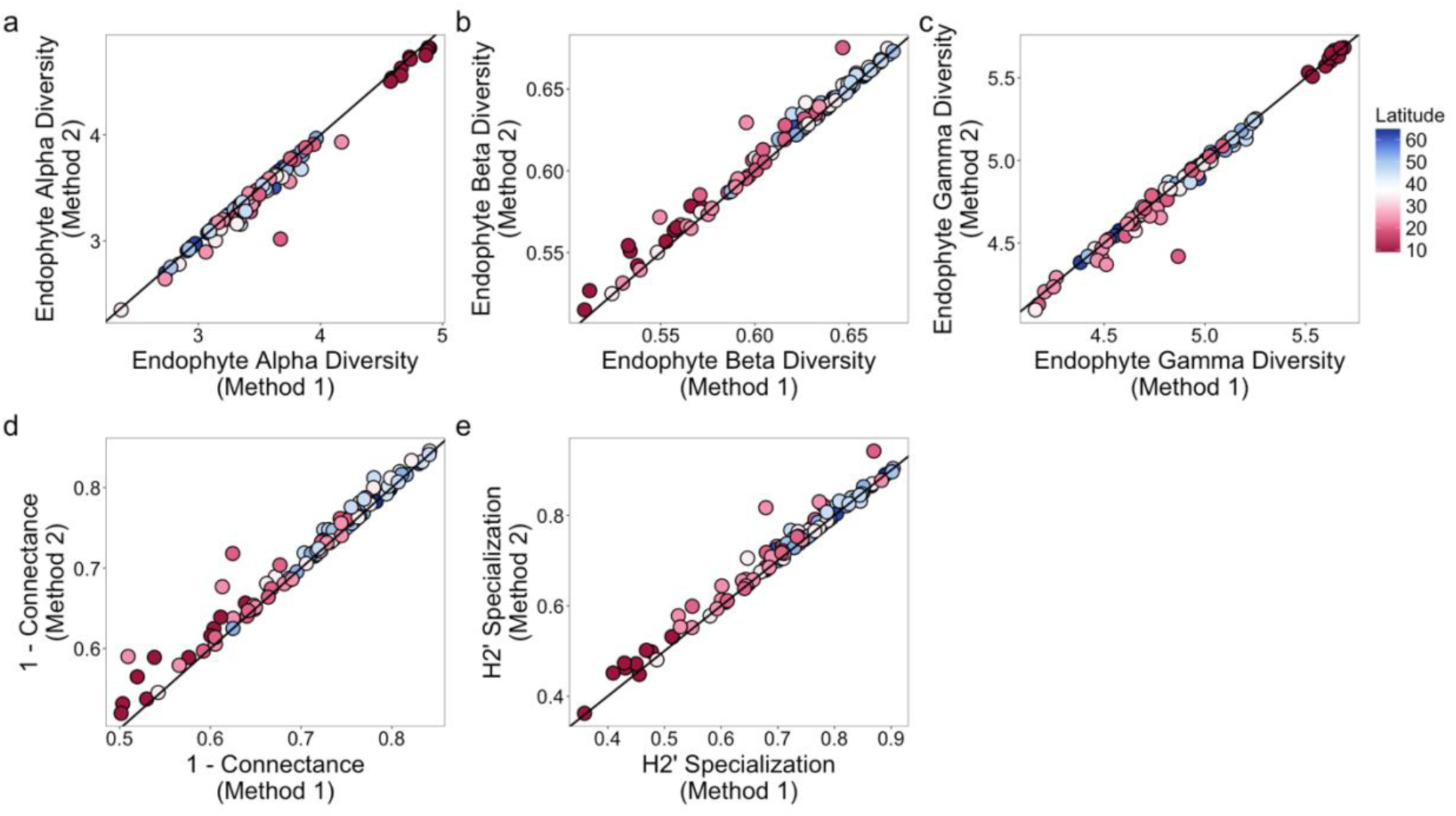
Comparison of methods on measures of community structure. Method 1 combined plant congeners within each plot by summing the read abundances between them, followed by rarefying. Method 2 randomly chose one representative plant congener in each plot to consider for the analyses and did not consider the other plant congeners. Diagonal line shown is a 1:1 line. Each point refers to one plot and colors refer to latitudinal position. (a) Endophyte alpha diversity (endophyte richness per plant sample) is highly correlated across the two methods (Pearson’s r: 0.99; p < 0.001). (b) Endophyte beta diversity (endophyte compositional similarity among host plants) is highly correlated across the two methods (Pearson’s r: 0.99; p < 0.001). (c) Endophyte gamma diversity (endophyte richness per plot) is highly correlated across the two methods (Pearson’s r: 0.99; p < 0.001). (d) Specialization measured as 1-connectance is highly correlated across the two methods (Pearson’s r: 0.98; p < 0.001). (e) Specialization measured as the H2’ index is highly correlated across the two methods (Pearson’s r: 0.98; p < 0.001).

**Figure S3.**
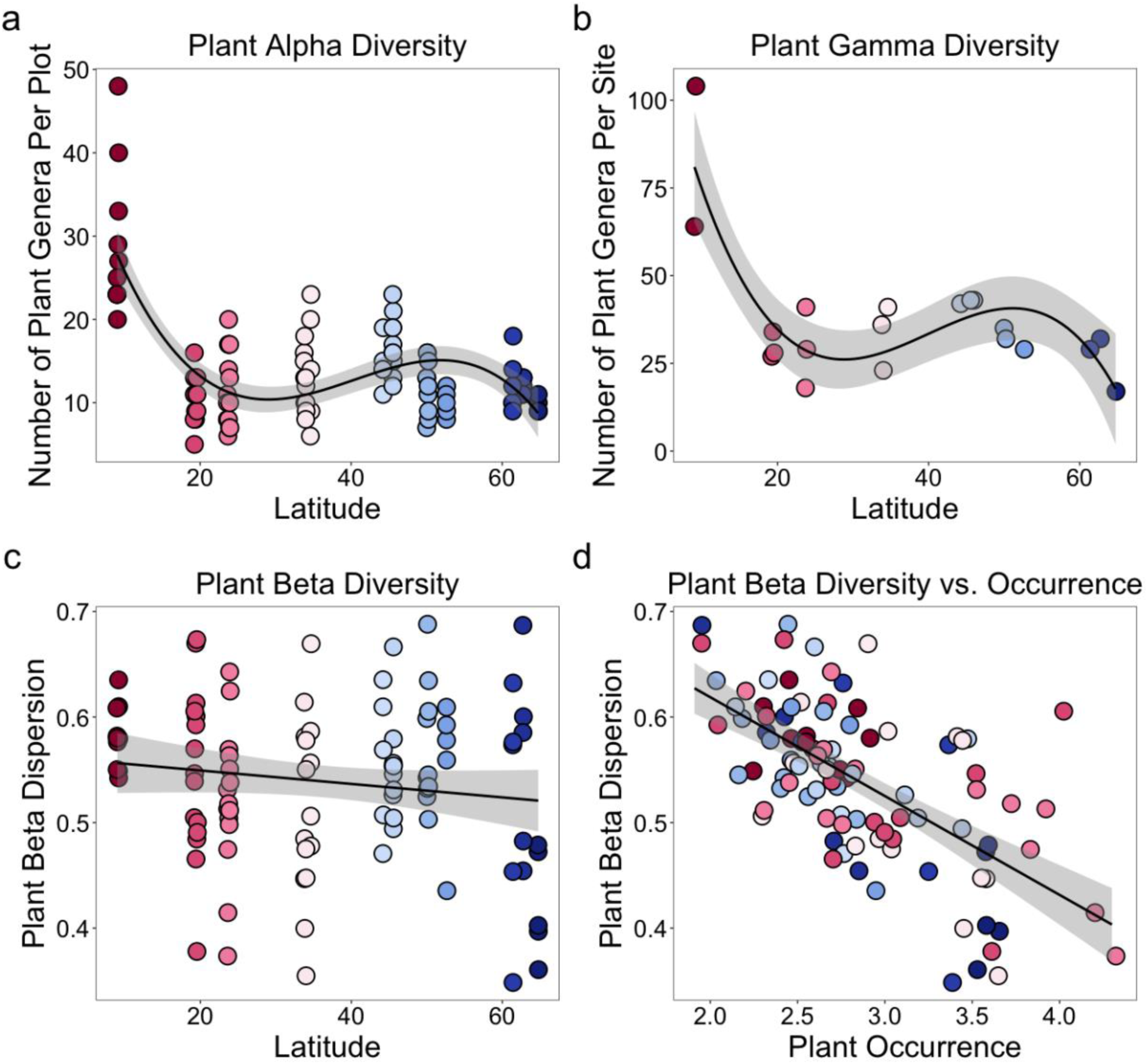
Relationships between measures of plant diversity and latitude (panels a-b), and the relationship between plant beta diversity and plant occurrence (panel d). (a, b) Plant alpha diversity (the number of plant genera in a plot) and plant gamma diversity (the number of plant genera in a site) are greatest in the tropical rainforests of Panama. (c) Plant beta diversity (the beta dispersion of plots in principal coordinate space relative to their site centroid) does not significantly vary across latitude. (d) Plant beta diversity is negatively correlated with plant occurrence: plots composed of plants found more frequently in a site had lower plant beta diversity (Pearson’s r: -0.64, p < 0.001).

**Figure S4.**
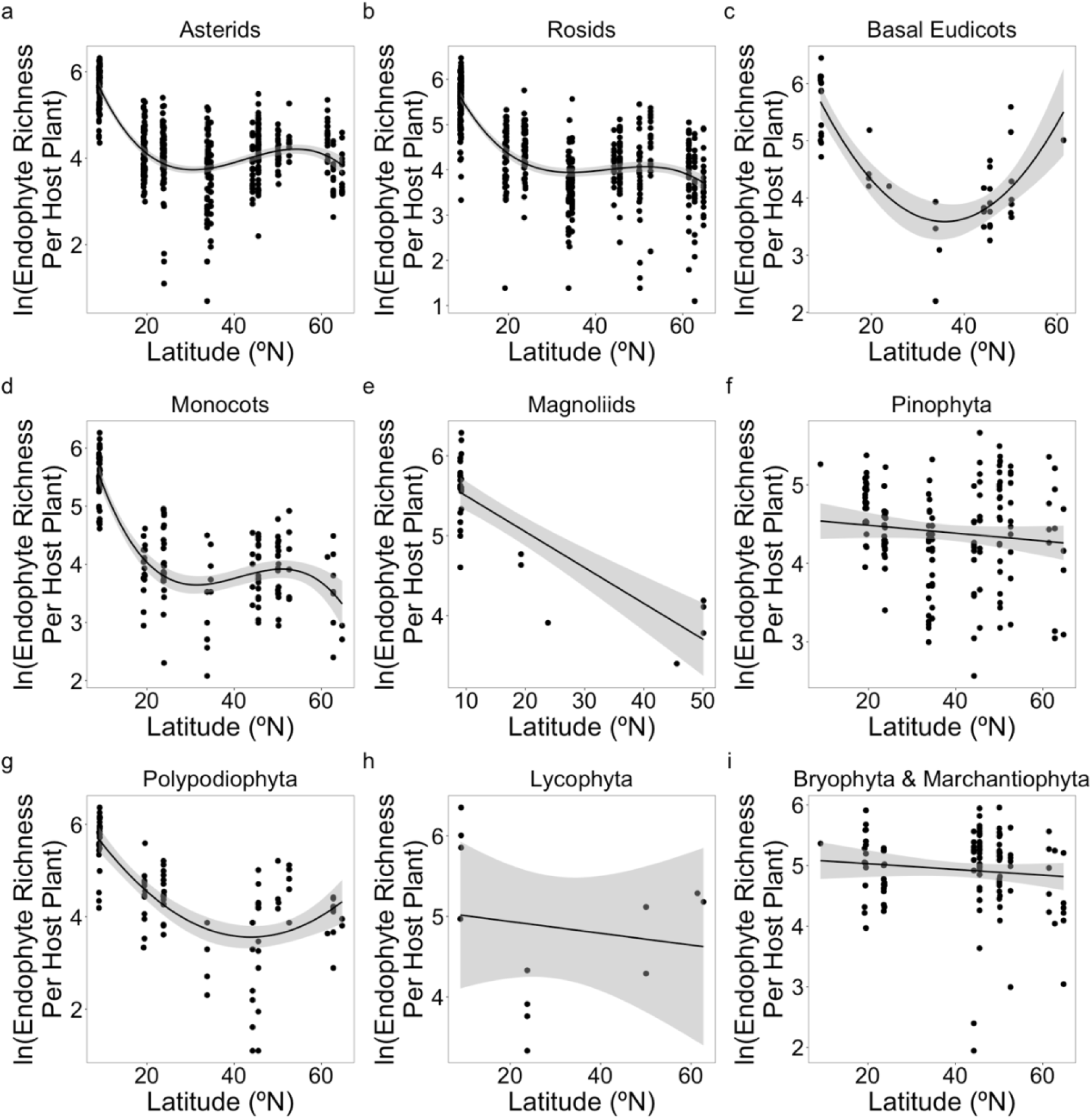
Endophyte richness as a function of latitude per major plant lineage. Each point is the 97% OTU richness of endophytes within a plant individual and is natural log scaled. Shaded regions are 95% confidence intervals. Plants were grouped into “major plant lineages” (Sessa et al. 2014, Li et al. 2021) as (a) asterids, (b) rosids, (c) basal eudicots, (d) monocots, (e) Magnoliids, (f) Pinophyta, (g) Polypodiophyta, (h) Lycophyta, and (i) Bryophyta or Marchantiophyta. Nonvascular plants (panel i) were removed for all analyses (see Methods).

**Figure S5.**
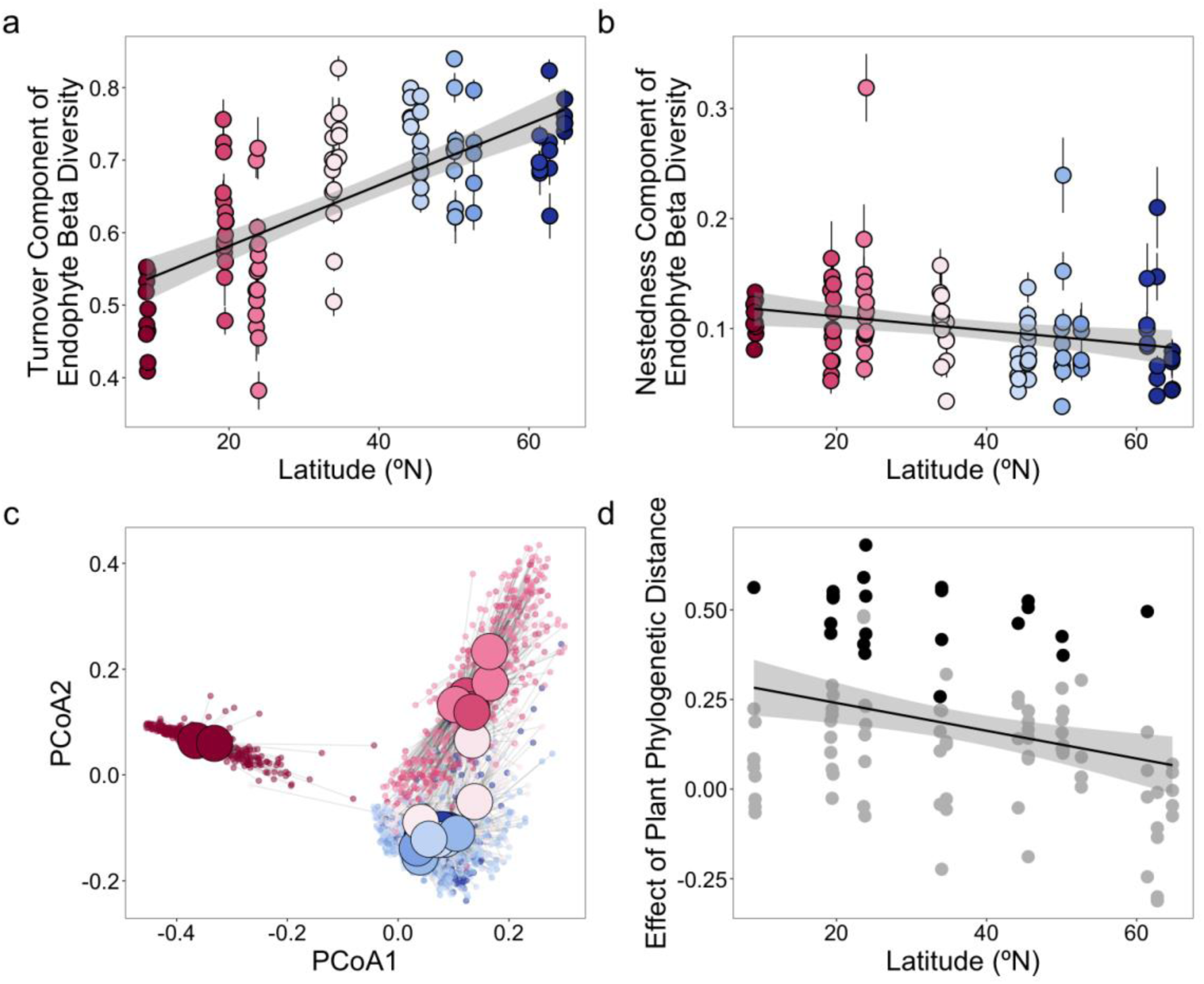
Decreasing endophyte beta diversity toward the tropics is driven by turnover in composition (panel a), rather than nestedness (panel b). The composition of endophyte communities is strongly influenced by site (panel c) and weakly influenced by plant phylogenetic distance (panel d). (a) The turnover component of endophyte beta diversity decreases toward the tropics (linear latitude term: p = 0.0001). (b) The nestedness component of endophyte beta diversity does not significantly vary across latitude (linear latitude term: p = 0.035). (c) Endophyte community composition (Bray-Curtis dissimilarity) in principal coordinate space. Small circles refer to the endophyte community associated with each plant individual (n = 1,539), and large circles refer to the centroid among endophyte communities within a given site (n = 20). (d) The effect of plant phylogenetic distance on endophyte composition (Mantel r) minimally increased toward the tropics latitudes when considering all plots (R^2^ = 0.084, p = 0.0019; solid line). Each point is a plot. Black points refer to plots with p-values < 0.05 and grey points refer to plots with p-values ≥ 0.05. Colors in panels a-c refer to latitudinal position.

**Figure S6.**
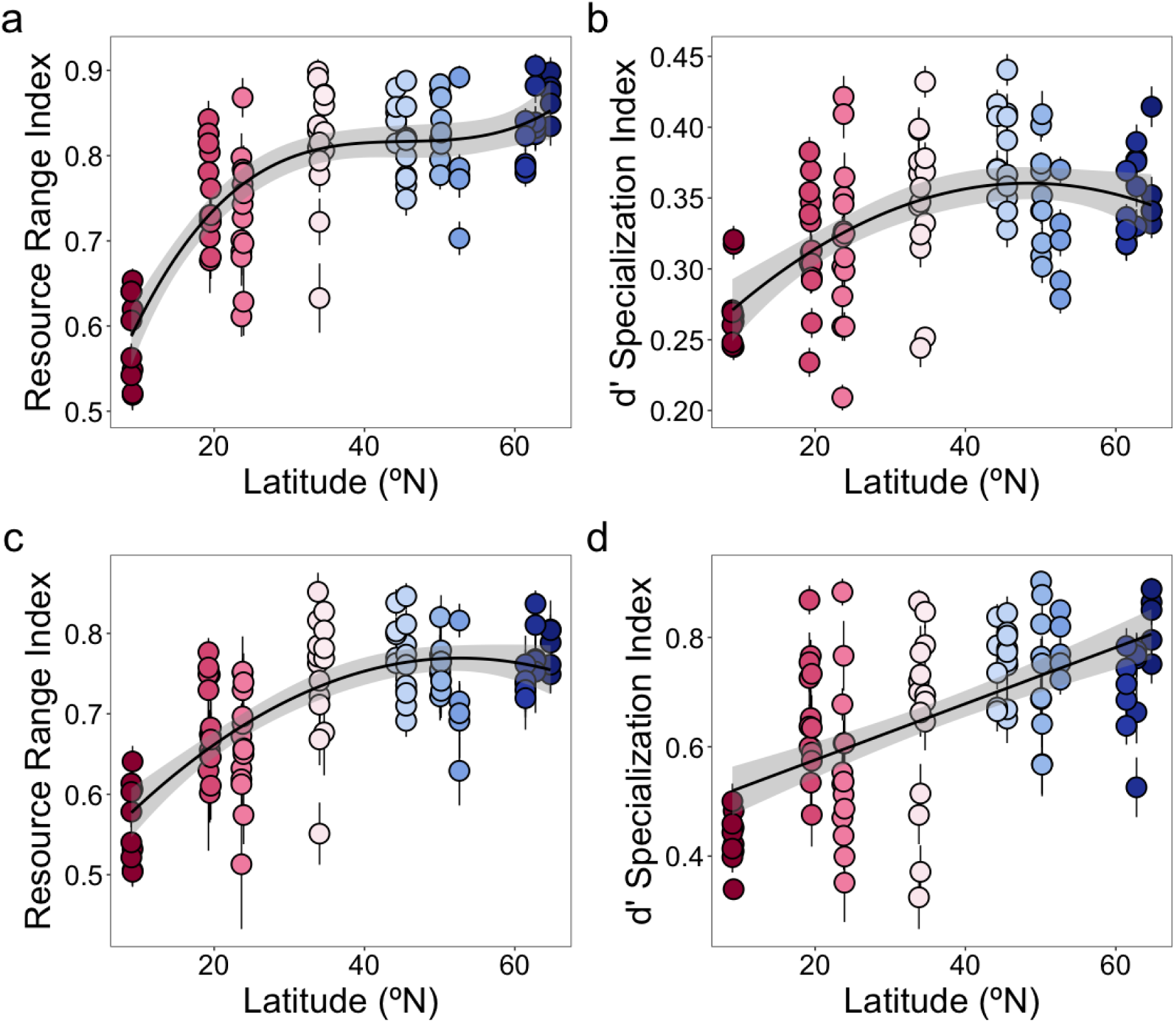
Alternative metrics of specialization that measure the specificity of endophytes to plants (panels a and b) or the specificity of plants to endophytes (panels c and d) decrease toward the tropics. (a) Endophyte resource range index decreases toward the tropics (R^2^ = 0.61; cubic latitude term: p = 0.036). (b) Endophyte d’ specialization index decreases toward the tropics (R^2^ = 0.29; quadratic latitude term: p = 0.011). (c) Plant resource range index decreases toward the tropics (R^2^ = 0.51; quadratic latitude term: p = 0.0061). (d) Plant d’ specialization index decreases toward the tropics (R^2^ = 0.35; linear latitude term: p < 0.001).

**Figure S7.**
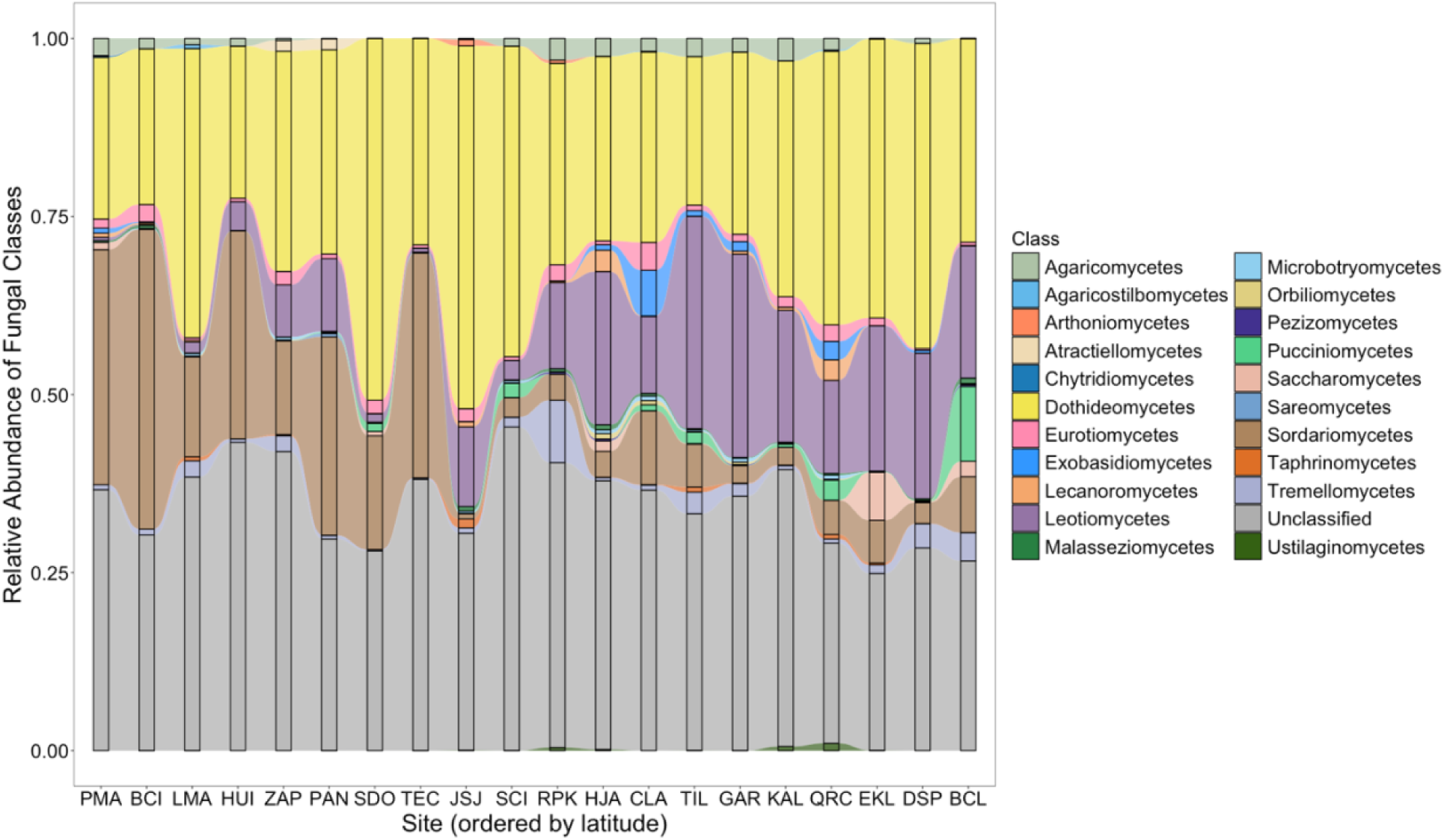
Relative abundance of fungal classes at the site level. Fungal endophyte OTUs are grouped by taxonomic class. Sites are ordered by latitudinal position. Sites PMA and BCI are located in Panamá Oeste, Panama. Sites LMA, HUI, and ZAP are located in Tlaxcala, Mexico. Site PAN is located in Durango, Mexico. Sites SDO and TEC are located in Zacatecas, Mexico. Sites JSJ, SCI, and RPK are located in California, USA. Sites HJA, CLA, and TIL are located in Oregon, USA. Sites GAR, KAL, and QRC are located in British Columbia, Canada. Sites EKL, DSP, and BCL are located in Alaska, USA.

**Figure S8.**
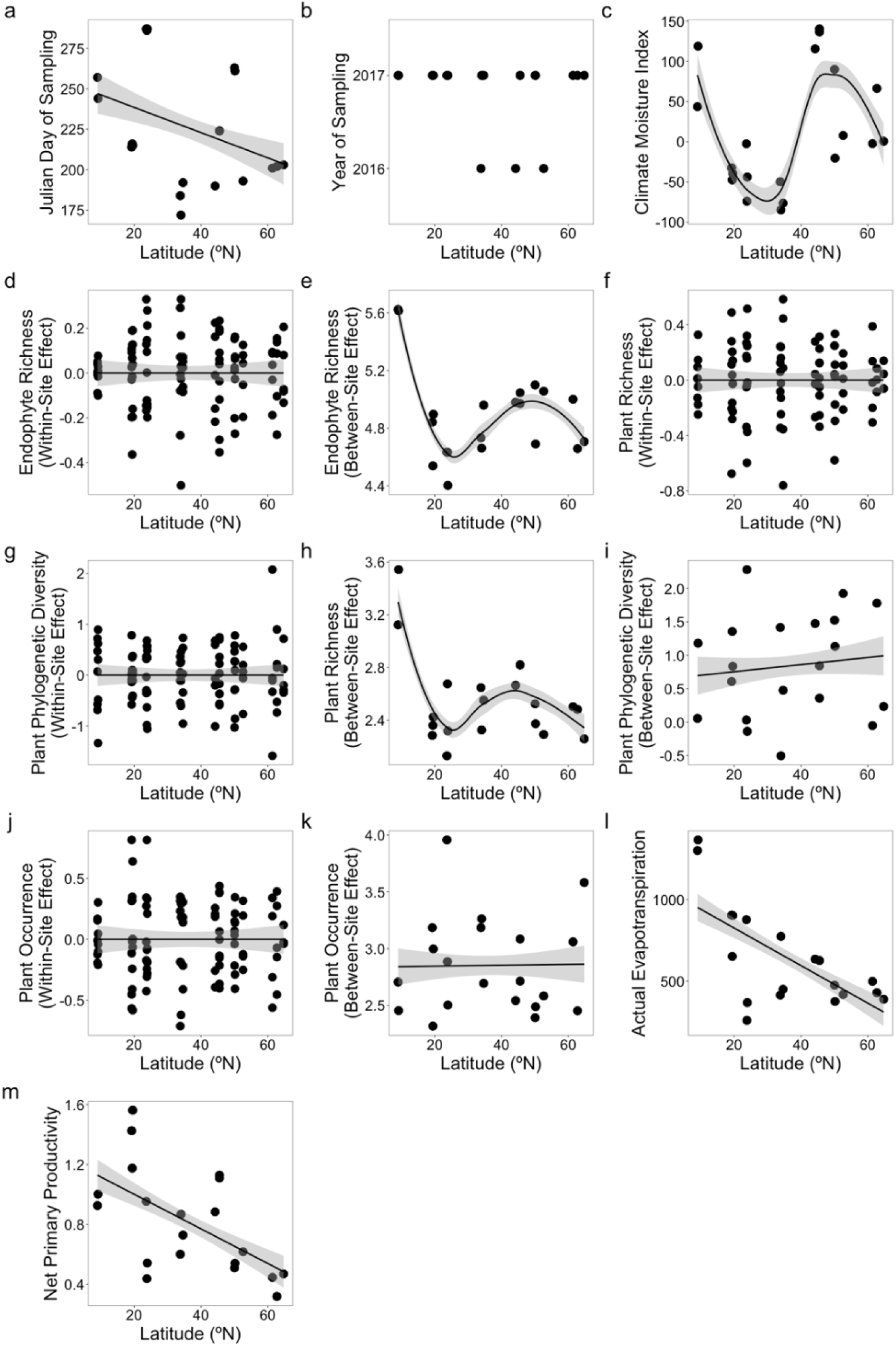
Covariates in the structural equation model. Each point represents one site (panels a, b, c, e, h, i, k, l, m) or one plot (panels d, f, g, j). Climate moisture index is measured in kg/m^2^/month, actual evapotranspiration is measured in kg/m²/year, and net primary productivity is measured in kg C/m²/year.

**Figure S9.**
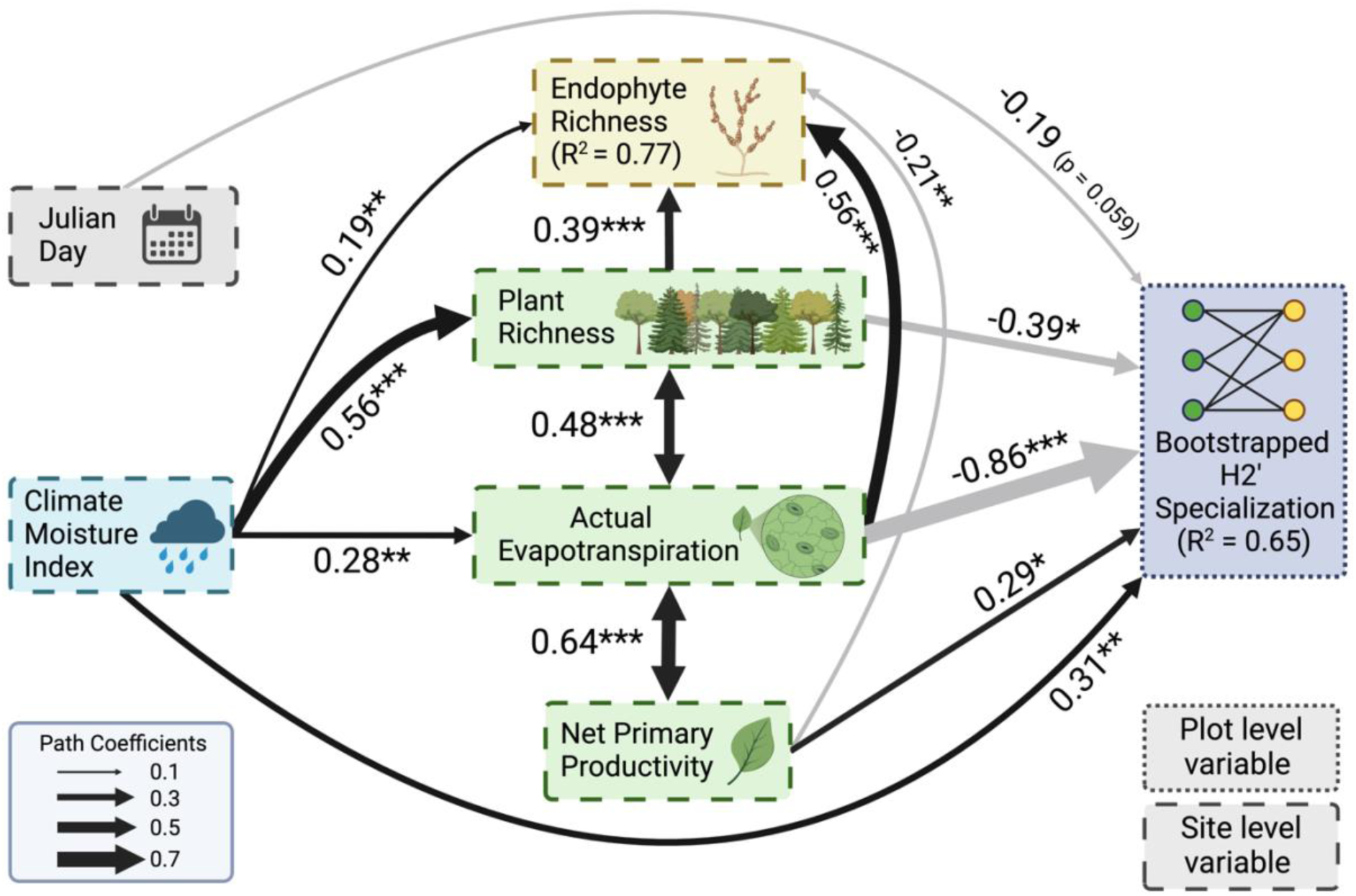
Piecewise structural equation model results using a bootstrapped H2’ index. Black and grey paths distinguish positive and negative relationships, respectively, in the structural equation model (Fisher’s C = 59.37, df = 80, p = 0.96). Unidirectional paths represent relationships in a simple or multiple linear regression, while bidirectional arrows represent correlated errors between variables. The width of each path corresponds to the magnitude of the standardized regression coefficient or the correlation coefficient. Paths shown are labeled with their statistical significance (***: 0 < p < 0.001, **: 0.001 < p < 0.01, *: 0.01 < p < 0.05) or their exact p-value if p > 0.05. R-squared values are the marginal r-squared values from linear mixed effect models. Boxes with dashed or solid lines are within- or between-site effects, respectively.

**Figure S10.**
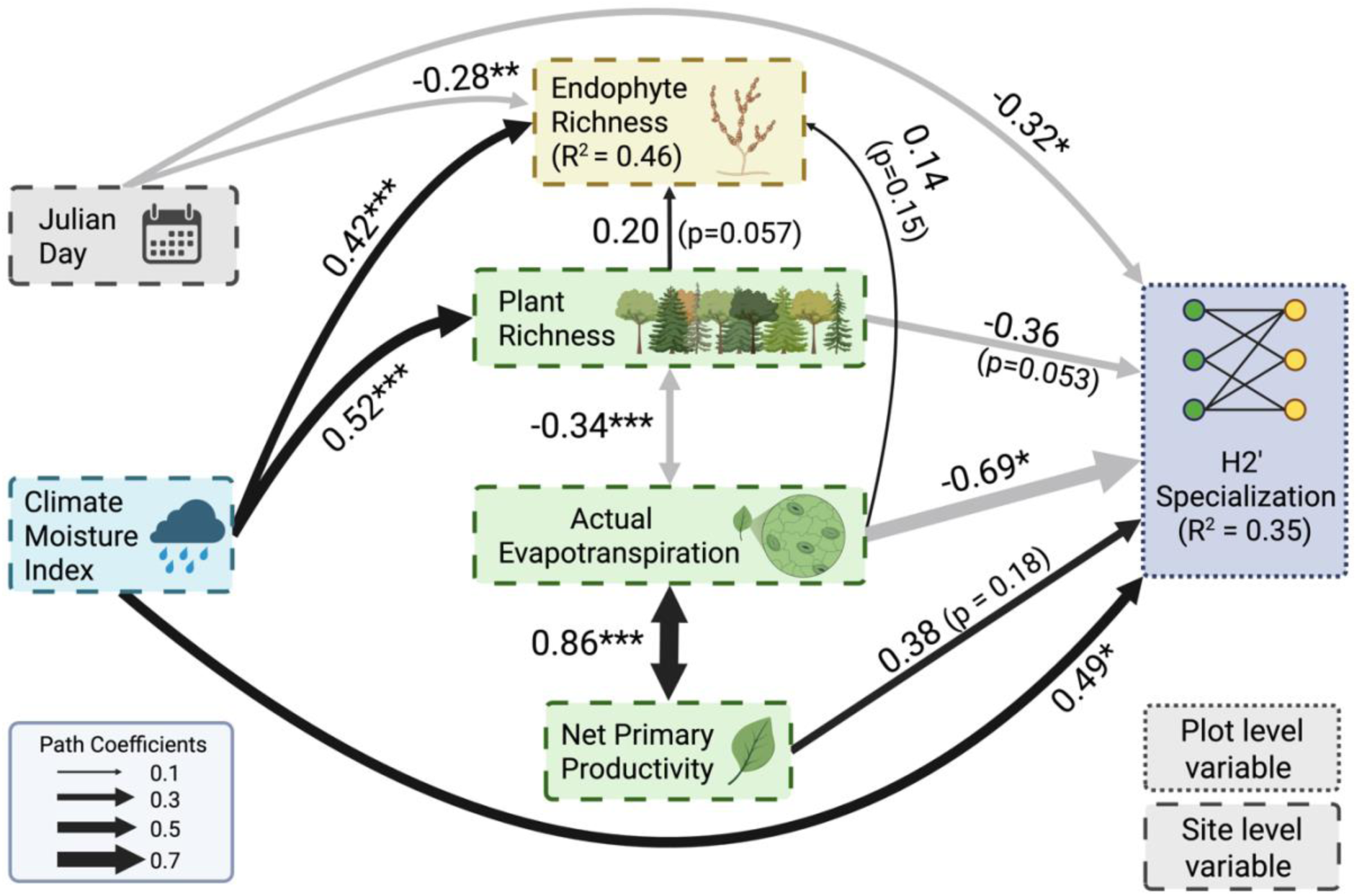
Piecewise structural equation model without the two tropical rainforest sites in Panama. Black and grey paths distinguish positive and negative relationships, respectively, in the structural equation model (Fisher’s C = 74.86, df = 72, p = 0.39). Unidirectional paths represent relationships in a simple or multiple linear regression, while bidirectional arrows represent correlated errors between variables. The width of each path corresponds to the magnitude of the standardized regression coefficient or the correlation coefficient. Paths shown are labeled with their statistical significance (***: 0 < p < 0.001, **: 0.001 < p < 0.01, *: 0.01 < p < 0.05) or their exact p-value if p > 0.05. R-squared values are the marginal r-squared values from linear mixed effect models. Boxes with dashed or solid lines are within- or between-site effects, respectively.

