## Supplementary Figures for "Specialization between foliar fungal endophyte and plant communities decreases toward the tropics"

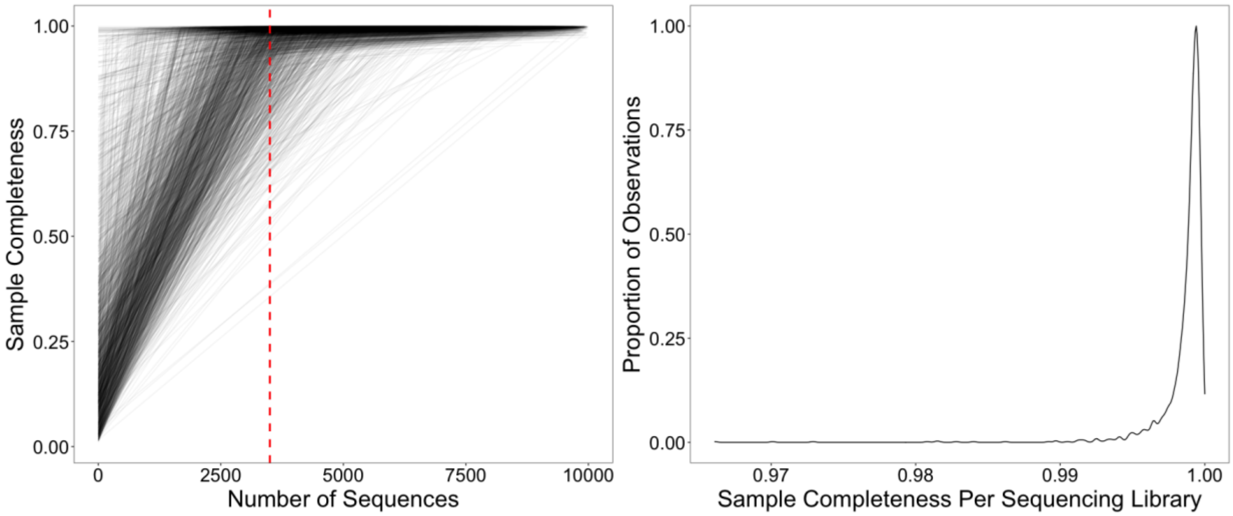


**Figure S1. Sample completeness, a metric for how well an endophyte community was characterized by sequencing, in relation to (panel a) sequencing depth or as a (panel b) histogram**. (a) Endophyte communities (each black line) were rarefied to a sequencing depth of 3500 reads (vertical red dashed line) because sample completeness tended to asymptote near that threshold. (b) The majority of the endophyte communities had sample completeness >99%.

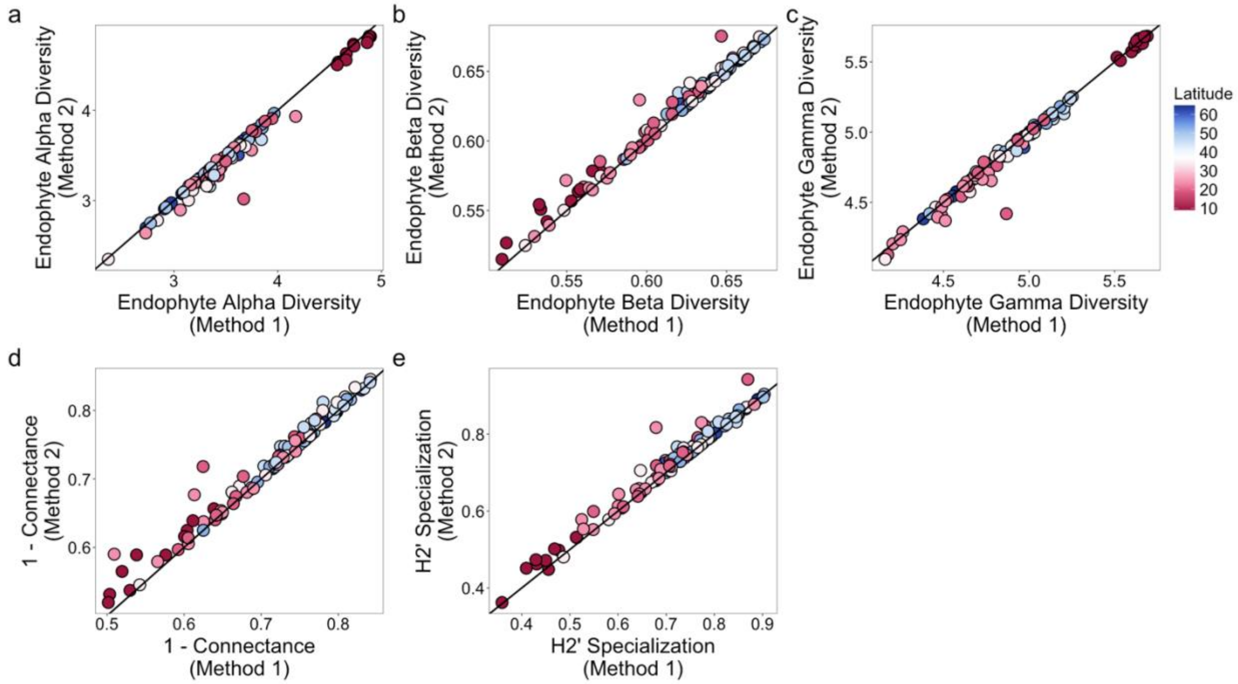


**Figure S2. Comparison of methods on measures of community structure.** Method 1 combined plant congeners within each plot by summing the read abundances between them, followed by rarefying. Method 2 randomly chose one representative plant congener in each plot to consider for the analyses and did not consider the other plant congeners. Diagonal line shown is a 1:1 line. Each point refers to one plot and colors refer to latitudinal position. (a) Endophyte alpha diversity (endophyte richness per plant sample) is highly correlated across the two methods (Pearson's r: 0.99; $p < 0.001$). (b) Endophyte beta diversity (endophyte compositional similarity among host plants) is highly correlated across the two methods (Pearson's r: 0.99; $p < 0.001$). (c) Endophyte gamma diversity (endophyte richness per plot) is highly correlated across the two methods (Pearson's r: 0.99; $p < 0.001$). (d) Specialization measured as 1-connectance is highly correlated across the two methods (Pearson's r: 0.98; $p < 0.001$). (e) Specialization measured as the H2' index is highly correlated across the two methods (Pearson's r: 0.98; $p < 0.001$).

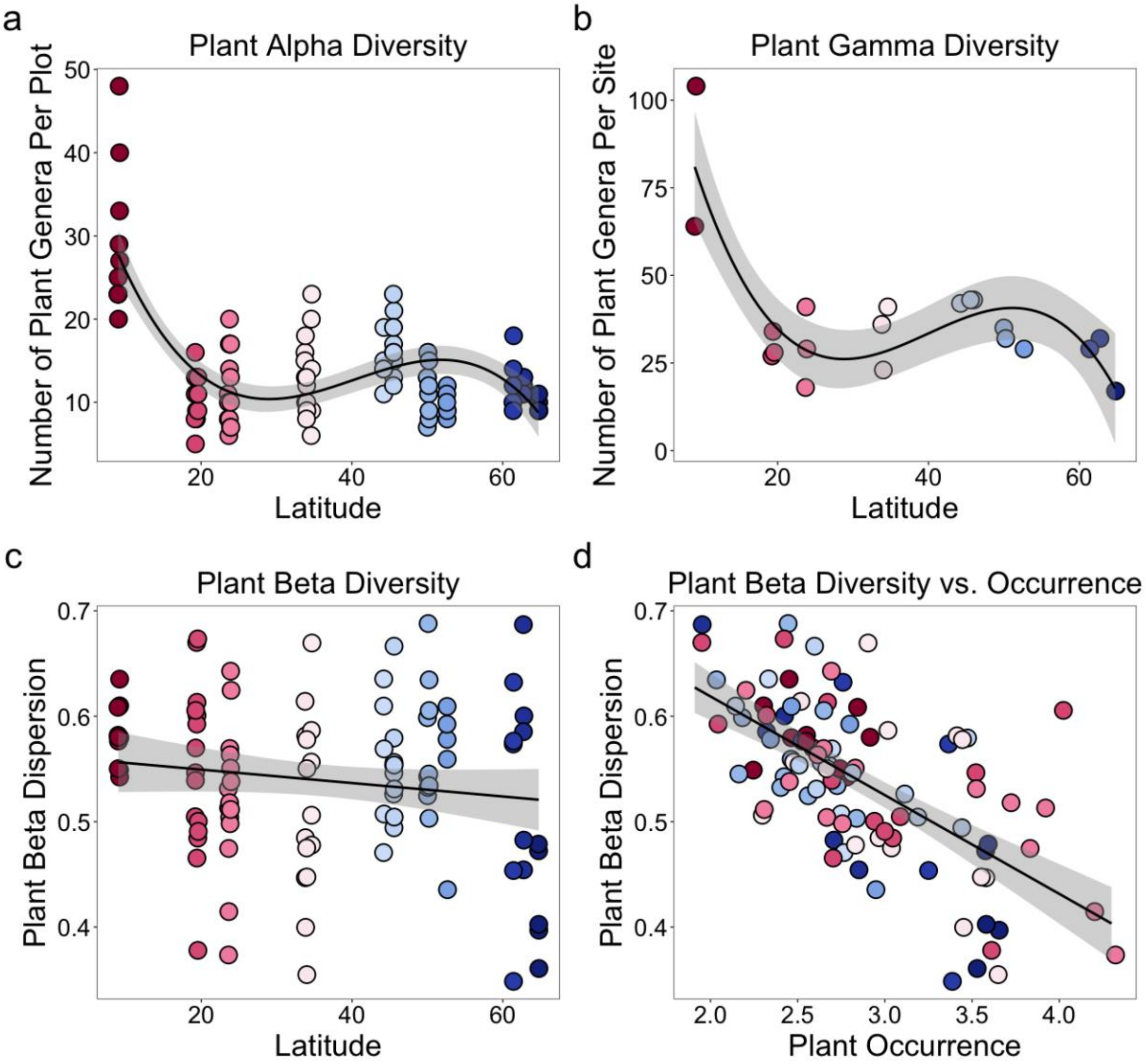


**Figure S3. Relationships between measures of plant diversity and latitude (panels a-b), and the relationship between plant beta diversity and plant occurrence (panel d).** (a, b) Plant alpha diversity (the number of plant genera in a plot) and plant gamma diversity (the number of plant genera in a site) are greatest in the tropical rainforests of Panama. (c) Plant beta diversity (the beta dispersion of plots in principal coordinate space relative to their site centroid) does not significantly vary across latitude. (d) Plant beta diversity is negatively correlated with plant occurrence: plots composed of plants found more frequently in a site had lower plant beta diversity (Pearson's r: -0.64, $p < 0.001$).

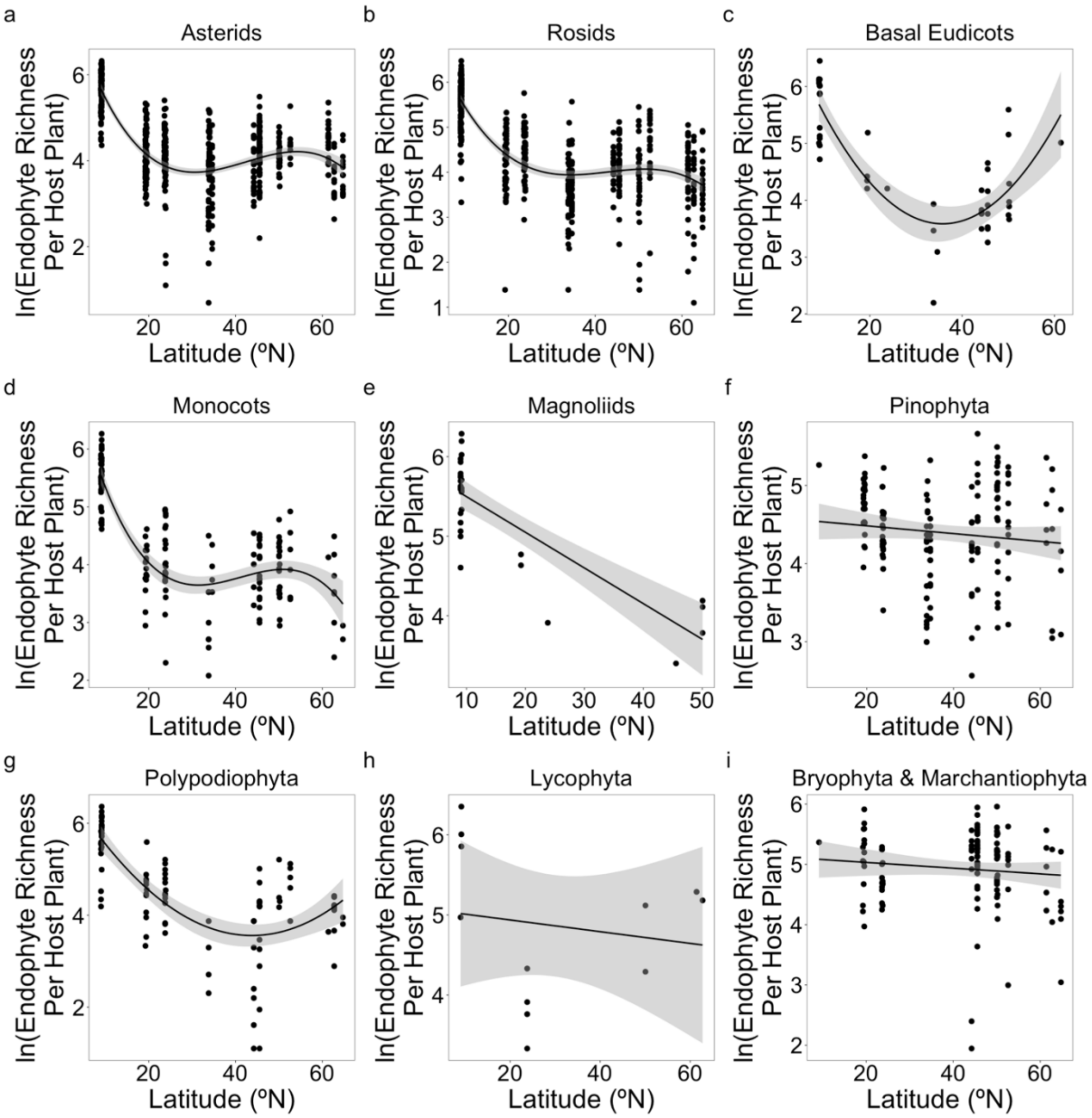


**Figure S4. Endophyte richness as a function of latitude per major plant lineage.** Each point is the 97% OTU richness of endophytes within a plant individual and is natural log scaled. Shaded regions are 95% confidence intervals. Plants were grouped into "major plant lineages" (Sessa et al. 2014, Li et al. 2021) as (a) asterids, (b) rosids, (c) basal eudicots, (d) monocots, (e) Magnoliids, (f) Pinophyta, (g) Polypodiophyta, (h) Lycophyta, and (i) Bryophyta or Marchantiophyta. Nonvascular plants (panel i) were removed for all analyses (see Methods).

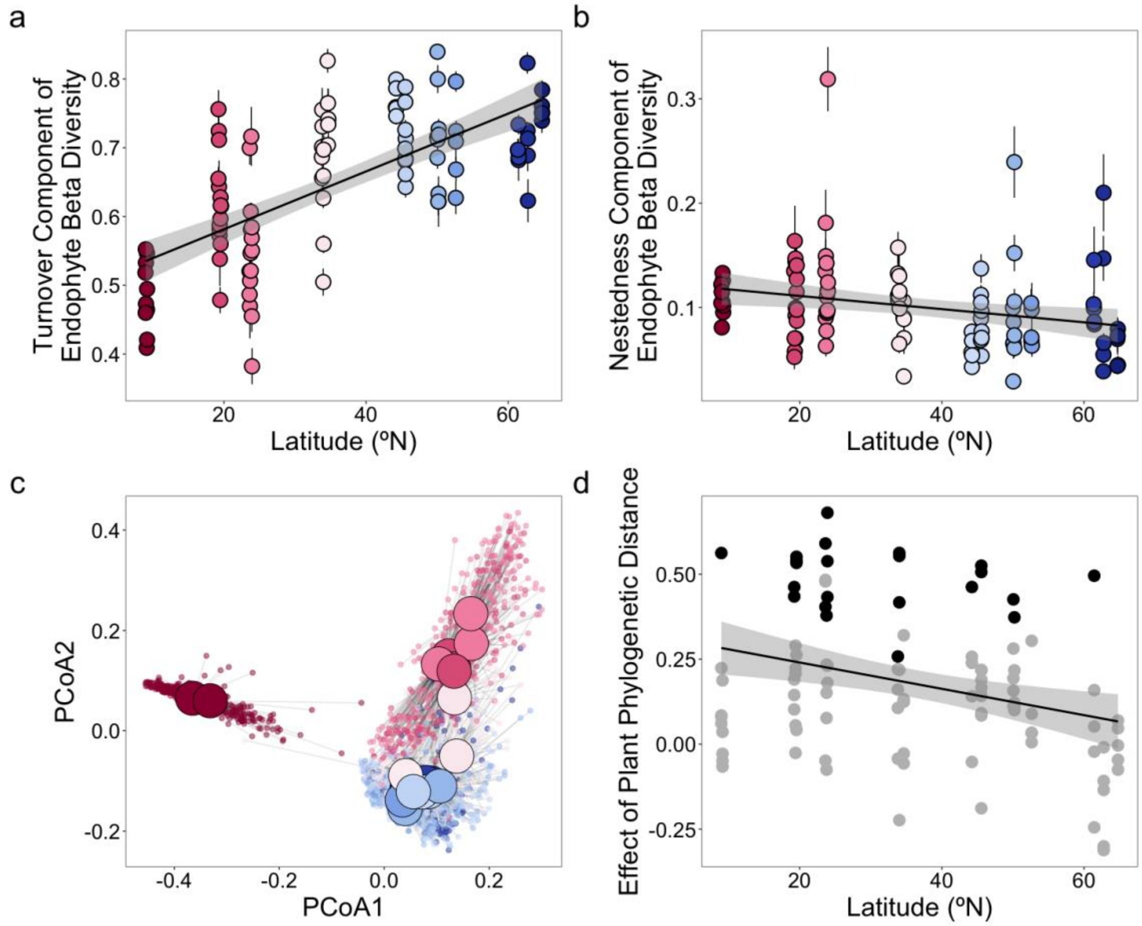


**Figure S5. Decreasing endophyte beta diversity toward the tropics is driven by turnover in composition (panel a), rather than nestedness (panel b). The composition of endophyte communities is strongly influenced by site (panel c) and weakly influenced by plant phylogenetic distance (panel d).** (a) The turnover component of endophyte beta diversity decreases toward the tropics (linear latitude term: p = 0.0001). (b) The nestedness component of endophyte beta diversity does not significantly vary across latitude (linear latitude term: p = 0.035). (c) Endophyte community composition (Bray-Curtis dissimilarity) in principal coordinate space. Small circles refer to the endophyte community associated with each plant individual (n = 1,539), and large circles refer to the centroid among endophyte communities within a given site (n = 20). (d) The effect of plant phylogenetic distance on endophyte composition (Mantel r) minimally increased toward the tropics latitudes when considering all plots ($R^2$ = 0.084, p = 0.0019; solid line). Each point is a plot. Black points refer to plots with p-values < 0.05 and grey points refer to plots with p-values ≥ 0.05. Colors in panels a-c refer to latitudinal position.

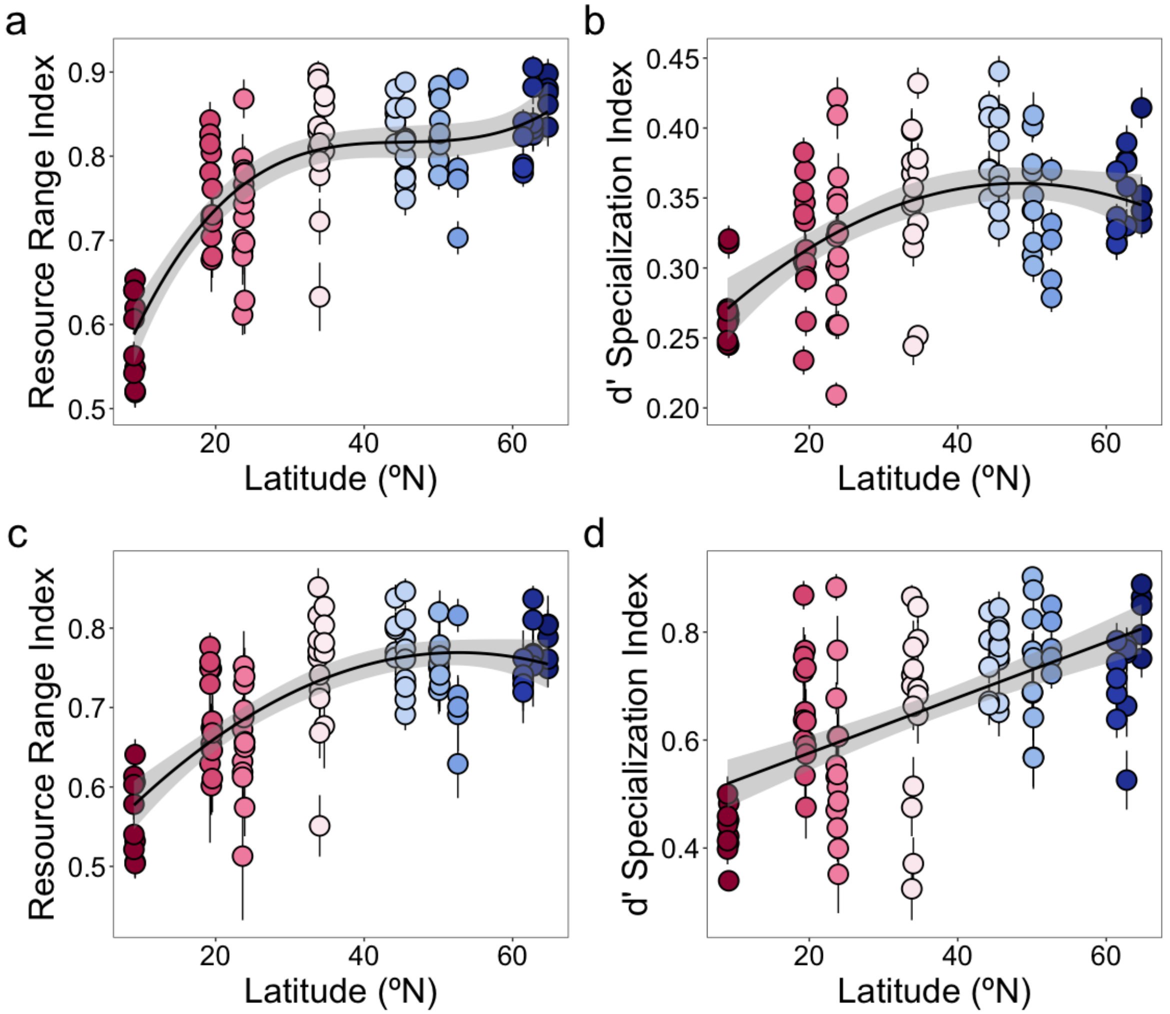


**Figure S6. Alternative metrics of specialization that measure the specificity of endophytes to plants (panels a and b) or the specificity of plants to endophytes (panels c and d) decrease toward the tropics.** (a) Endophyte resource range index decreases toward the tropics ($R^2$ = 0.61; cubic latitude term: $p$ = 0.036). (b) Endophyte d' specialization index decreases toward the tropics ($R^2$ = 0.29; quadratic latitude term: $p$ = 0.011). (c) Plant resource range index decreases toward the tropics ($R^2$ = 0.51; quadratic latitude term: $p$ = 0.0061). (d) Plant d' specialization index decreases toward the tropics ($R^2$ = 0.35; linear latitude term: $p < 0.001$).

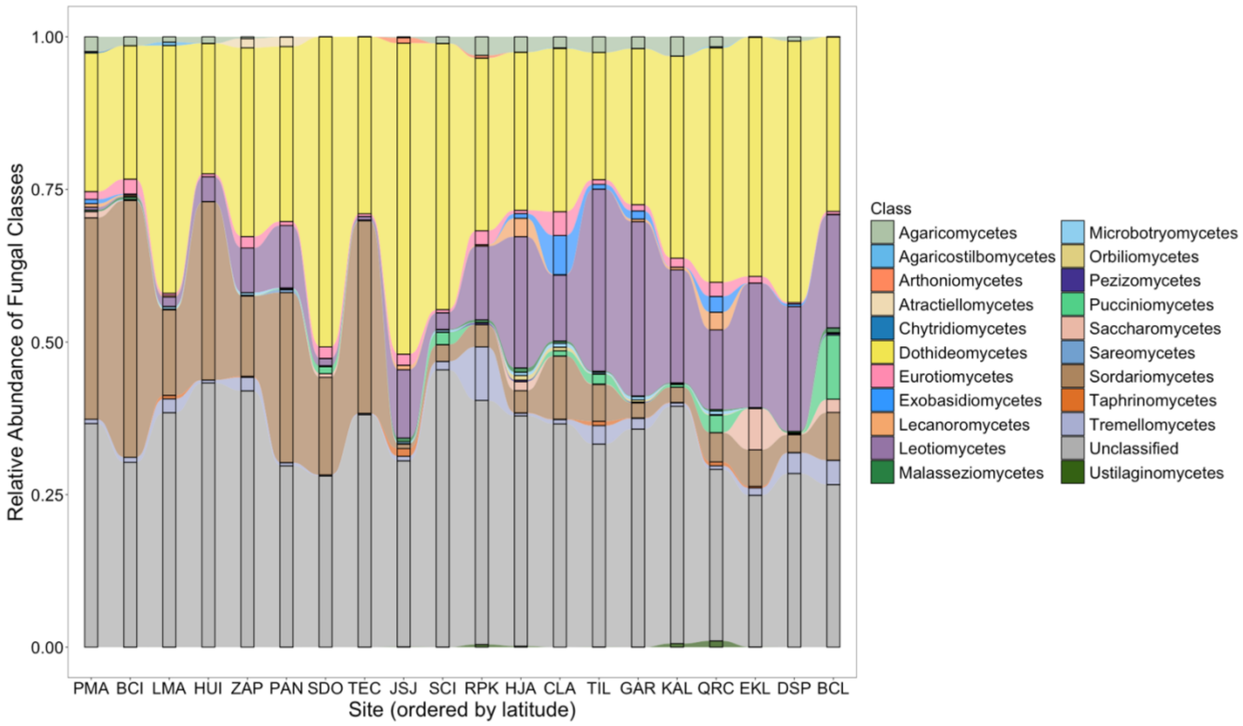


**Figure S7. Relative abundance of fungal classes at the site level.** Fungal endophyte OTUs are grouped by taxonomic class. Sites are ordered by latitudinal position. Sites PMA and BCI are located in Panamá Oeste, Panama. Sites LMA, HUI, and ZAP are located in Tlaxcala, Mexico. Site PAN is located in Durango, Mexico. Sites SDO and TEC are located in Zacatecas, Mexico. Sites JSJ, SCI, and RPK are located in California, USA. Sites HJA, CLA, and TIL are located in Oregon, USA. Sites GAR, KAL, and QRC are located in British Columbia, Canada. Sites EKL, DSP, and BCL are located in Alaska, USA.

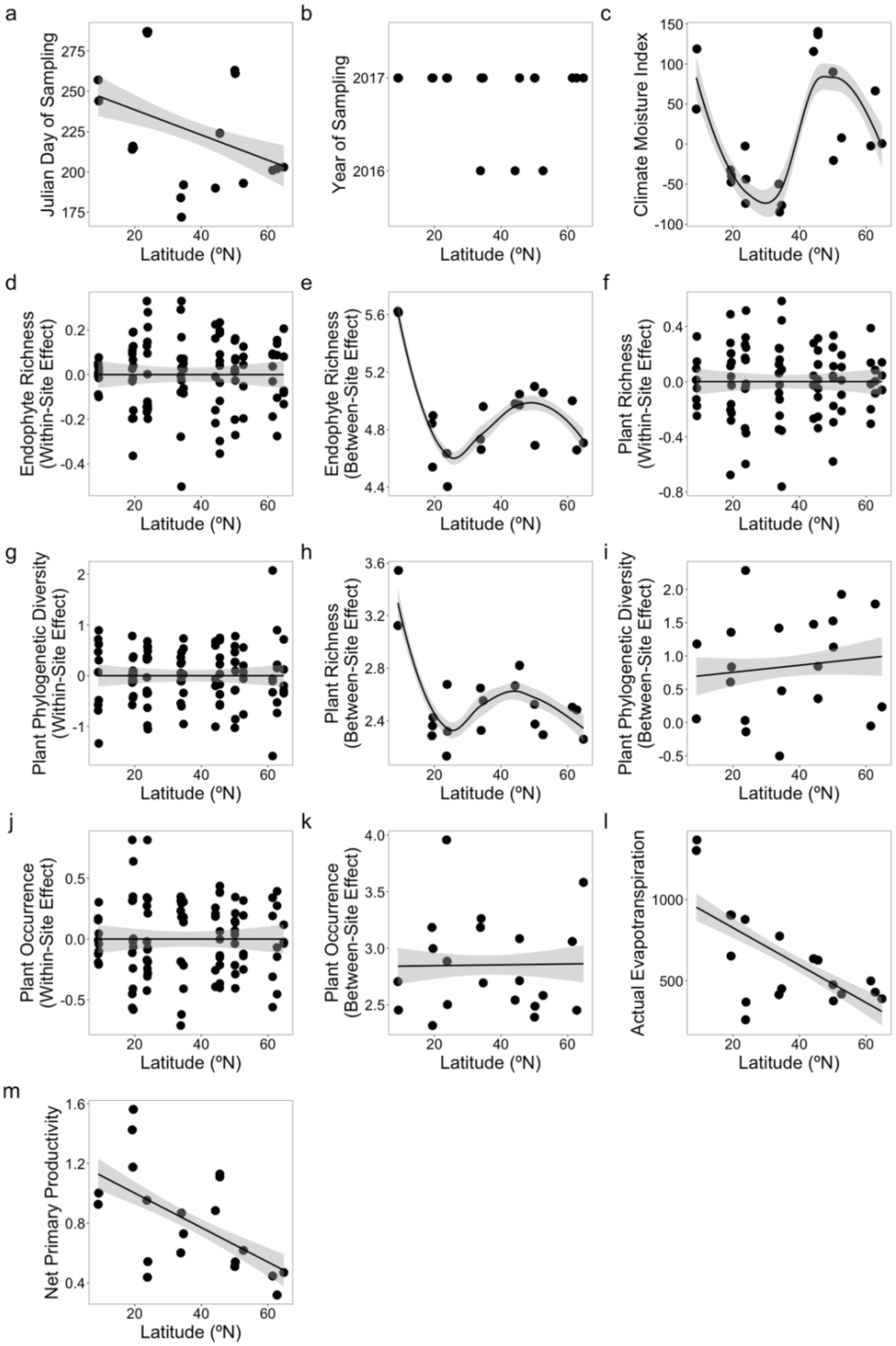


**Figure S8. Covariates in the structural equation model.** Each point represents one site (panels a, b, c, e, h, i, k, l, m) or one plot (panels d, f, g, j). Climate moisture index is measured in kg/m$^2$/month, actual evapotranspiration is measured in kg/m²/year, and net primary productivity is measured in kg C/m²/year.

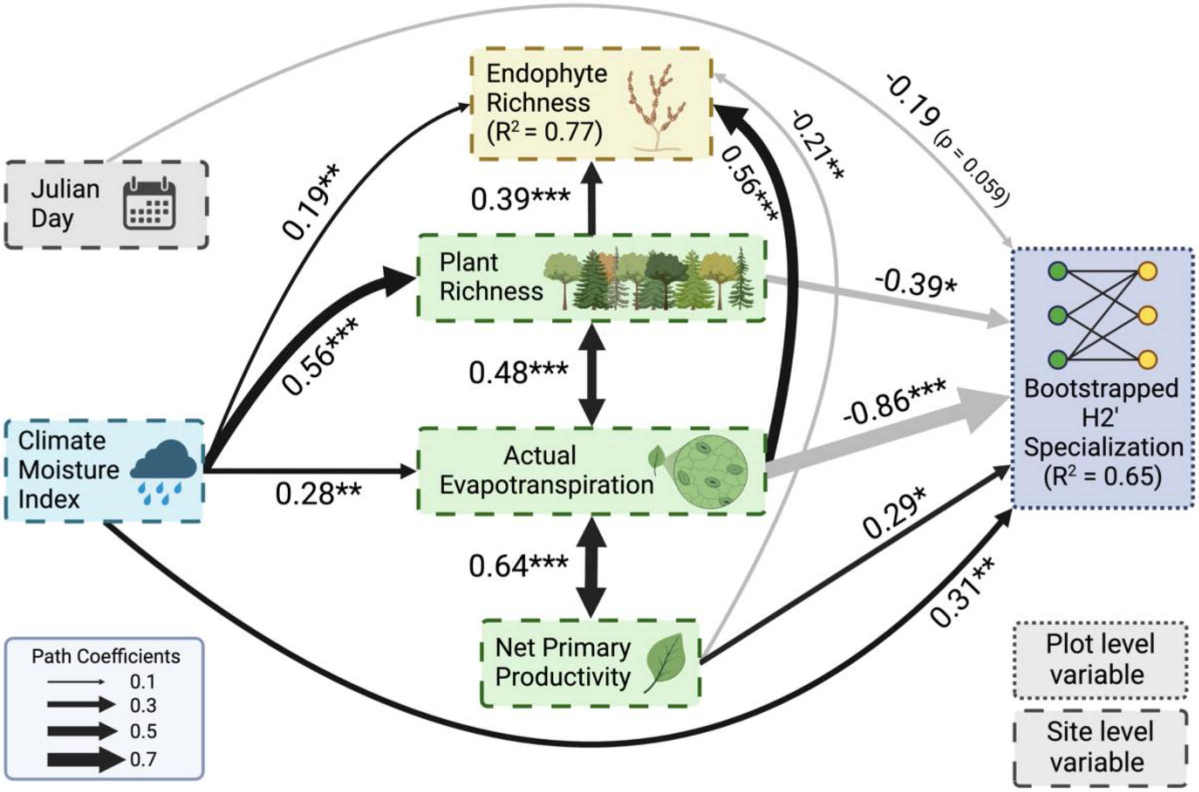


**Figure S9. Piecewise structural equation model results using a bootstrapped H2' index.** Black and grey paths distinguish positive and negative relationships, respectively, in the structural equation model (Fisher's C = 59.37, df = 80, p = 0.96). Unidirectional paths represent relationships in a simple or multiple linear regression, while bidirectional arrows represent correlated errors between variables. The width of each path corresponds to the magnitude of the standardized regression coefficient or the correlation coefficient. Paths shown are labeled with their statistical significance (***: $0 < p < 0.001$, **: $0.001 < p < 0.01$, *: $0.01 < p < 0.05$) or their exact p-value if $p > 0.05$. R-squared values are the marginal r-squared values from linear mixed effect models. Boxes with dashed or solid lines are within- or between-site effects, respectively.

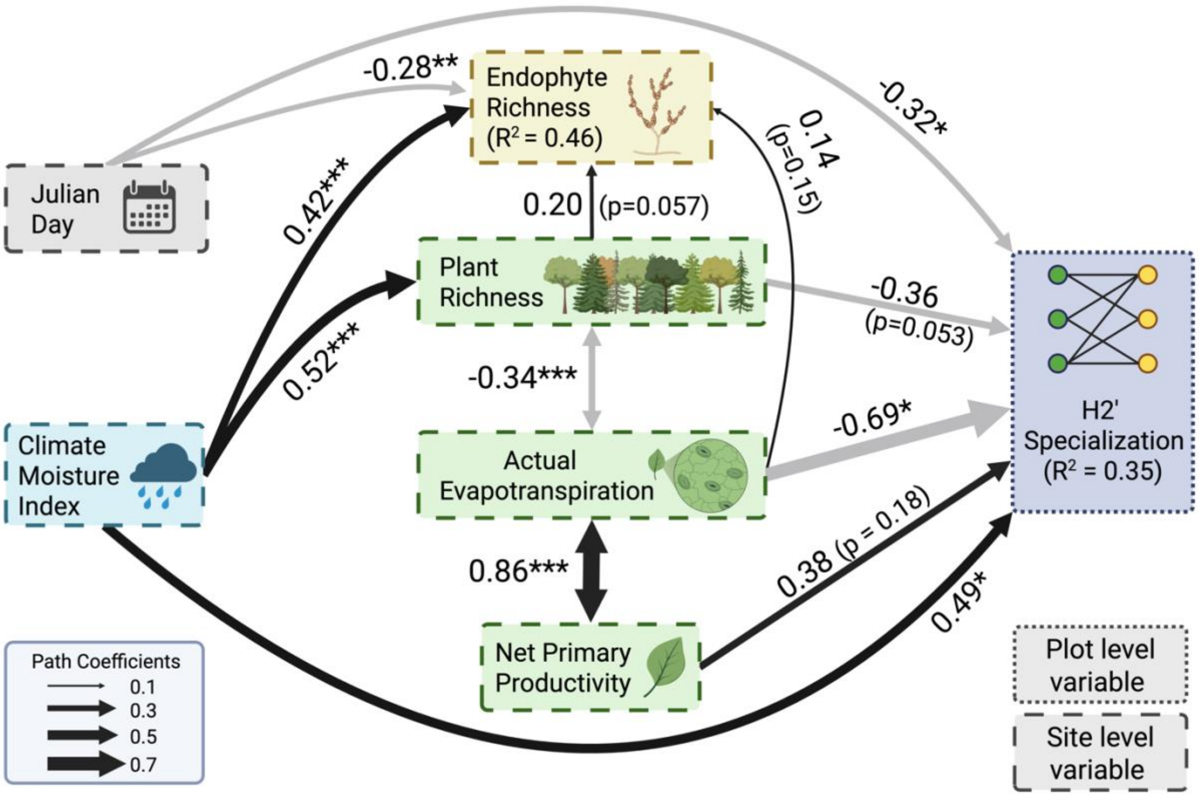


**Figure S10. Piecewise structural equation model without the two tropical rainforest sites in Panama.** Black and grey paths distinguish positive and negative relationships, respectively, in the structural equation model (Fisher's C = 74.86, df = 72, p = 0.39). Unidirectional paths represent relationships in a simple or multiple linear regression, while bidirectional arrows represent correlated errors between variables. The width of each path corresponds to the magnitude of the standardized regression coefficient or the correlation coefficient. Paths shown are labeled with their statistical significance (***: $0 < p < 0.001$, **: $0.001 < p < 0.01$, *: $0.01 < p < 0.05$) or their exact p-value if $p > 0.05$. R-squared values are the marginal r-squared values from linear mixed effect models. Boxes with dashed or solid lines are within- or between-site effects, respectively.
